# Molecular and Neural Circuit Mechanisms Underlying Sexual Experience-dependent Long-Term Memory in *Drosophila*

**DOI:** 10.1101/2024.09.28.615582

**Authors:** Dongyu Sun, Xiaoli Zhang, Hongyu Miao, Tianmu Zhang, Woo Jae Kim

## Abstract

The neural and molecular underpinnings of sexual experience-dependent long-term memory (SELTM) in male *Drosophila melanogaster* remain poorly understood, despite its significance for reproductive success. Here, we dissect the role of a specific class of neurons, termed ’Yuelao’ (YL) neurons, in the formation of SELTM and the associated molecular pathways. We employed a combination of genetic manipulations, immunohistochemistry, and behavioral assays in *Drosophila* to investigate the function of YL neurons and their molecular interactors in SELTM. Utilizing the RNA sequencing data and transgenic tools, we delineated the neural circuits involved in taste and pheromone processing relevant to SELTM. Our findings reveal that the YL neurons, expressing the Orb2 scaffolding protein, are indispensable for the formation of SELTM following sexual experience. These neurons are regulated by the neuromodulator short neuropeptide F (sNPF) and its receptor (sNPF-R), which modulate glutamate release via NMDAR2. We demonstrate that sexual experience triggers synaptic plasticity in YL neurons, characterized by an increase in dendritic and presynaptic terminal areas, and a decrease in intracellular calcium levels. Furthermore, we show that YL neurons are specialized for the generation of appetitive sexual experience-dependent memory and project to brain regions implicated in memory formation. Our study uncovers the YL neurons as a key neural substrate for SELTM in *Drosophila*, shedding light on the molecular and circuit mechanisms that mediate the formation of long-term memories following sexual experience. These findings provide novel insights into the neural basis of taste-related memory and have broader implications for understanding the interplay between experience, memory formation, and behavior.

## INTRODUCTION

*Drosophila melanogaster* is a widely used model organism for investigating the brain circuitry involved in memory formation and retrieval. The memory systems of the fruit fly, such as visual, olfactory, and taste, have been well studied and each has specific neural substrates^1^. The mushroom body (MB), a crucial component of the *Drosophila* brain, predominantly facilitates the process of olfactory and visual learning and memory^2^. Olfactory memory is dependent on the antennal lobe (AL), which is the convergence point for olfactory receptor neurons. Additionally, the MB α’/β’ lobes play a role in olfactory memory by receiving inputs from the AL and processing olfactory information ^3^. Visual memory relies on the optic lobes, which receive visual inputs, and the central complexes (CC) including the ellipsoid body (EB) and the fan-shaped body (FB)^4–6^. In addition, the dorsal accessory calyx (dAC), which is composed of the a/b Kenyon cells, primarily receives visual input^7–9^.

Despite the extensive knowledge accumulated on these sensory memories, the neural mechanisms underlying sexual experience-dependent long-term-memory (SELTM) in male *Drosophila* remain relatively uncharted. Unlike olfactory memory, which has been extensively dissected at both the molecular and circuit levels, SELTM is less explored, and its neural underpinnings are not as clearly defined. This gap in understanding is particularly notable given the significance of SELTM in reproductive success and species survival^10^. Sexual experience can induce long-lasting changes in male courtship behavior^11^, which indicates that this behavioral adaptation is underpinned by associative learning and memory processes. However, the specific neural pathways and molecular cascades that mediate this type of memory are not well understood.

While experience-dependent long-term-memory (LTM) has been well established in the context of courtship conditioning in *Drosophila*^12,13^, the precise mechanisms by which actual sexual experience shapes memory traces remain enigmatic. Courtship conditioning, a form of associative learning, has been extensively studied and involves the modification of male courtship behavior based on prior experience with females^12,14–18^. However, the intricate ways in which the act of mating itself can sculpt the neural architecture and memory formation in males are not well understood.

A particularly intriguing aspect of sexual experience in *Drosophila* is the dichotomy between the well-documented post-mating behaviors of females^19^, such as reduced receptivity and increased egg laying^20–25^, and the relatively unexplored realm of male sexual experience and its impact on memory and future behaviors^26,27^. Unlike females, where the neurobiological changes of post-mating have been linked to specific physiological and behavioral adaptations, the memory-related consequences of sexual experience in males are less characterized. The scarcity of research in this area is surprising, considering the potential insights SELTM could provide into the evolution of reproductive strategies and the complexity of memory processing in the brain. This knowledge could also have broader implications for understanding the neural basis of complex behaviors and the interplay between sensory experiences and memory consolidation^28^.

The *orb2* gene encodes a postsynaptic scaffolding protein that is indispensable for the establishment of experience-dependent long-term courtship memory in *Drosophila*^29^. This protein has the capacity to form amyloid-like oligomers, a process that alters its molecular function, transitioning from a translation repressor to an activator^30^. Orb2 can facilitate memory formation through two distinct mechanisms: by binding to target mRNA 3’ untranslated regions (UTRs) to augment the translation of genes involved in neuronal growth, synapse formation, and protein turnover^31,32^ or independently of its RNA-binding affinity^30,33–35^. Although the role of Orb2 in mediating olfactory and courtship memory in *Drosophila* has been well-established, its involvement in SELTM remains to be fully elucidated.

This study aims to bridge this gap by investigating the neural and molecular signatures of sexual experience in male *Drosophila*, offering novel insights into how such experiences are encoded and retrieved within the brain. By delineating the function of Orb2-expressing MB projection neurons, which are situated outside the conventional MB Kenyon cell population, we aim to find the molecular mechanisms modulating memory processes that are specifically associated with gustatory and pheromonal sensory inputs to SELTM.

## RESULTS

### Sexual experience induces Orb2-specific LTM in male *Drosophila*

We previously elucidated that taste and pheromonal cues exert a pivotal influence on the modulation of mating investment by sexual experience in male *Drosophila*^36,37^. This behavior, termed ’shorter-mating-duration (SMD)’, is a form of interval timing that is prevalent across the animal kingdom, enabling organisms to quantify time intervals within the range of seconds to minutes^38^. The internal clock model posits that the coordination of pacemaker-accumulator circuits with memory circuits within the brain is essential for the precise comparison of time durations^39^. Consequently, we posited that the SMD behavior is contingent upon the engagement of memory circuits and associated traces within the neural architecture of the brain (Fig. S1A)^40–43^.

Males undergone sexual experience demonstrate a persistent reduction in MD, with this behavioral pattern persisting even 12 hours post-isolation from females (Fig. 1A-E), indicating the establishment of LTM. To corroborate that this attenuation of MD is a consequence of cumulative mating experiences, we conducted experiments on serial mating of males, which revealed that repetitive mating did not diminish mating duration (Fig. 1F-H). This finding suggests that the reduction in mating investment is contingent upon the formation of LTM.

**Fig 1.**
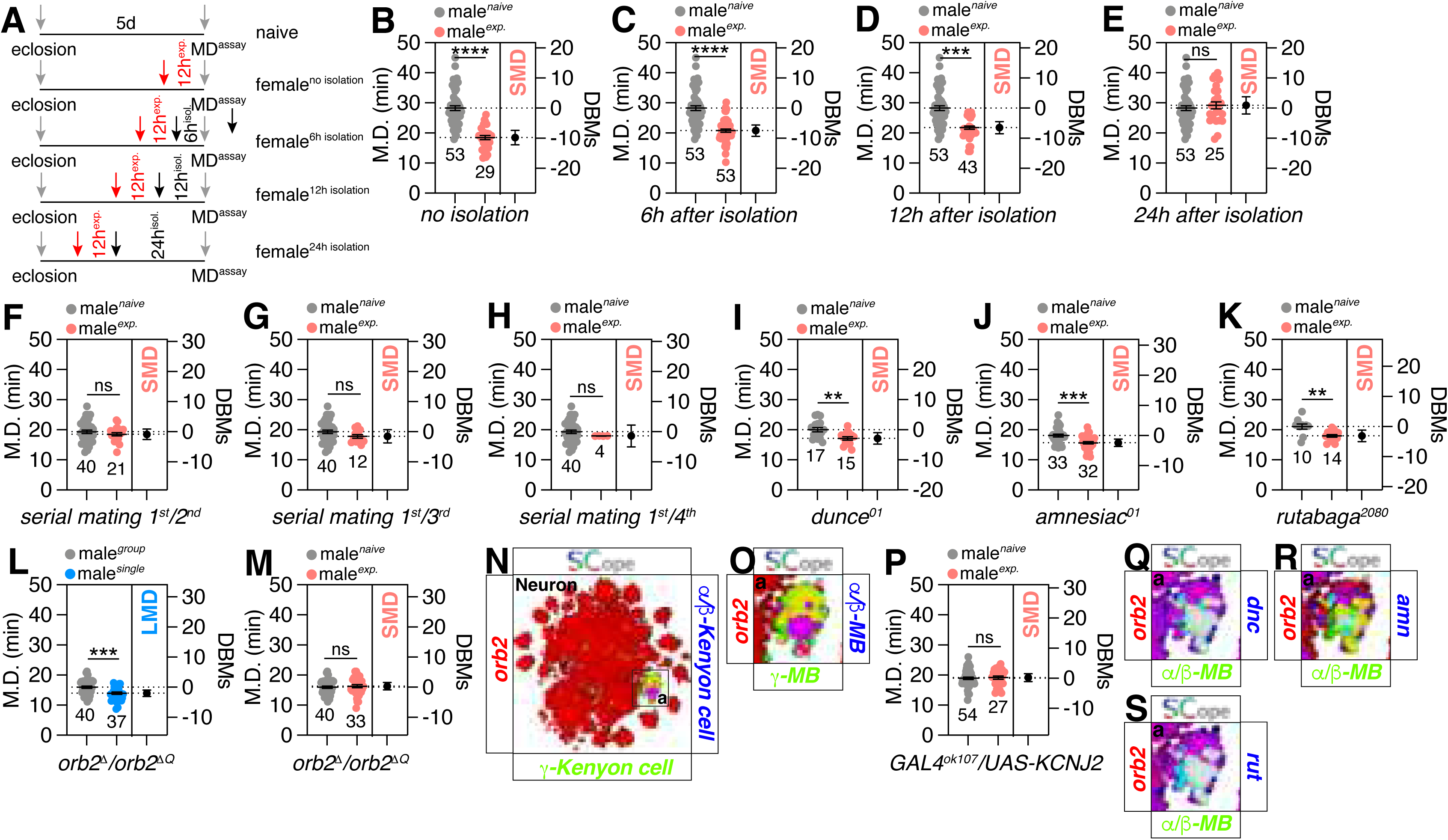
Sexual experience elicits a robust memory that endures for over 12 hours. (A) The sexual experience-isolation experiment. The red arrow indicates the starting time point for doubling the number of female fruit flies and males, and the black arrow indicates the time point for returning male fruit flies to sexual isolation. The duration of each phase is indicated in the figure. MD experiments have been conducted for all groups after a period of 5 days. The DBMs states that “Difference between means” in context of estimation statistics. The Difference between means refers to the disparity or variation in the average values of two distinct datasets or groups. (B-E) SMD assays for *Canton-S* after sexual isolation. The periods of isolation are as follows: no isolation (B), 6-hour isolation (C), 12-hour isolation (D), 24-hour isolation(E). Light grey dots represent naïve males and pink dots represent experienced males. Dot plots represent the MD of each male fly. The mean value and standard error are labeled within the dot plot (black lines). with error bars representing SEM. Asterisks represent significant differences, as revealed by the Student’s *t* test and ns represents non-significant difference (*\*p<0.05, **p<0.01, ***p< 0.001, ****p< 0.0001*). For detailed methods, see the METHODS for a detailed description of the mating duration assay used in this study. (F-H) Mating duration assays for *Canton-S,* naive (1^st^ mating) *Canton-S* VS exp. (2^nd^mating) *Canton-S* (F), naive (1^st^ mating) *Canton-S* VS exp. (3^rd^mating) *Canton-S* (G) and naive (1^st^ mating) *Canton-S* VS exp. (4^th^mating) *Canton-S* (H). (I-K) SMD assays for Memory-defective mutant fruit flies, *dunce^01^*(I)*, amnesiac^01^* (J) *and rutabaga*^2080^ (K). (L-M) LMD and SMD assays for Memory-defective mutant fruit flies *orb2^Δ^/orb2^ΔQ^* Light grey dots represent Group naïve males and blue dots represent single males. Dot plots represent the MD of each male fly.(N-O) Single-cell RNA sequencing (SCOPE scRNA-seq) datasets reveal cell clusters colored by expression of *orb2*(red) α/β-Kenyon cell (blue) and g-Kenyon cell (green) in brain. (P) SMD assays for *GAL4^ok107^* mediated electrical silencing *via UAS-KCNJ2*. (Q-S) Single-cell RNA sequencing (SCOPE scRNA-seq) datasets reveal cell clusters colored by expression of orb2 (red) dnc (blue) and α/β-Kenyon cell (α/β-MB, green) in neurons (Q). *orb2*(red) amn (blue) and α/β-Kenyon cell (α/β-MB, green) in neurons (R) and *orb2*(red) rut (blue) and α/β-Kenyon cell (α/β-MB, green) in neurons (S).

Previously, our group demonstrated that male fly prolongs their mating duration when confronted with competitors, a phenomenon termed as longer-mating-duration (LMD), which is also a type of interval timing behavior. This extended mating duration is contingent on the visual learning and memory capacity of the males, specifically linked to the expression of *amnesic* (*amn*) and *rutabaga* (*rut*) in specific EB rings (Fig. S1B-D)^44^.

Contrary to the LMD behavior, the LTM for SMD does not rely on the functions of *dnc*, *amn*, or *rut* (Fig. 1I-K). Instead, our data indicate that Orb2 is specifically implicated in SMD but not LMD (Fig. 1L-M). Analysis of the Fly SCope RNA sequencing dataset suggests that Orb2 is predominantly expressed in the α/β- and γ-lobe of the MB (Fig. 1N-O)^45^, aligning with previous reports (Fig. S1E) ^33^. Within the MB, which is a pivotal high-order brain region for memory processing, we have previously shown that inhibition of the EB but not the MB or FB disrupts LMD behavior (Fig. S1F-H)^44^. Conversely, inhibition of both the MB and EB, but not the FB, impairs SMD behavior (Fig. 1P and Fig. S1I-J). Furthermore, Fly SCope data predict that Orb2-expressing MB neurons also co-express *dnc*, *amn*, and *rut* (Fig. 1Q-S). In conjunction with genetic control data (Fig. S1K-N), the collective results indicate that LTM formation for SMD is contingent upon the functional engagement of the MB and EB, along with the function of the Orb2 gene in a currently undefined neuronal subset.

### The expression of Orb2 in a specific subset of MB projection neurons is pivotal for taste related LTM for generating SMD

The targeted knockdown of Orb2 in *GAL4^ok1^*^07^-positive MB neurons, but not in *GAL4^c5^*^47^-positive EB or *GAL4*^14–94^-positive FB neurons, disrupts SMD behavior (Fig. 2A-C). Utilizing advanced genetic tools designed to dissect MB circuits^46,47^, we conducted a screening to identify MB neurons that require Orb2 expression for the generation of SELTM (Table 1). Remarkably, our analysis has determined that the expression of Orb2 within neurons marked by *GAL4^MB093C^* is uniquely essential for the manifestation of SMD behavior (Fig. 2D, Fig. S2A-B for genetic control experiments, and Fig. S2C for independent *orb2-RNAi* strain confirmation).

**Fig 2.**
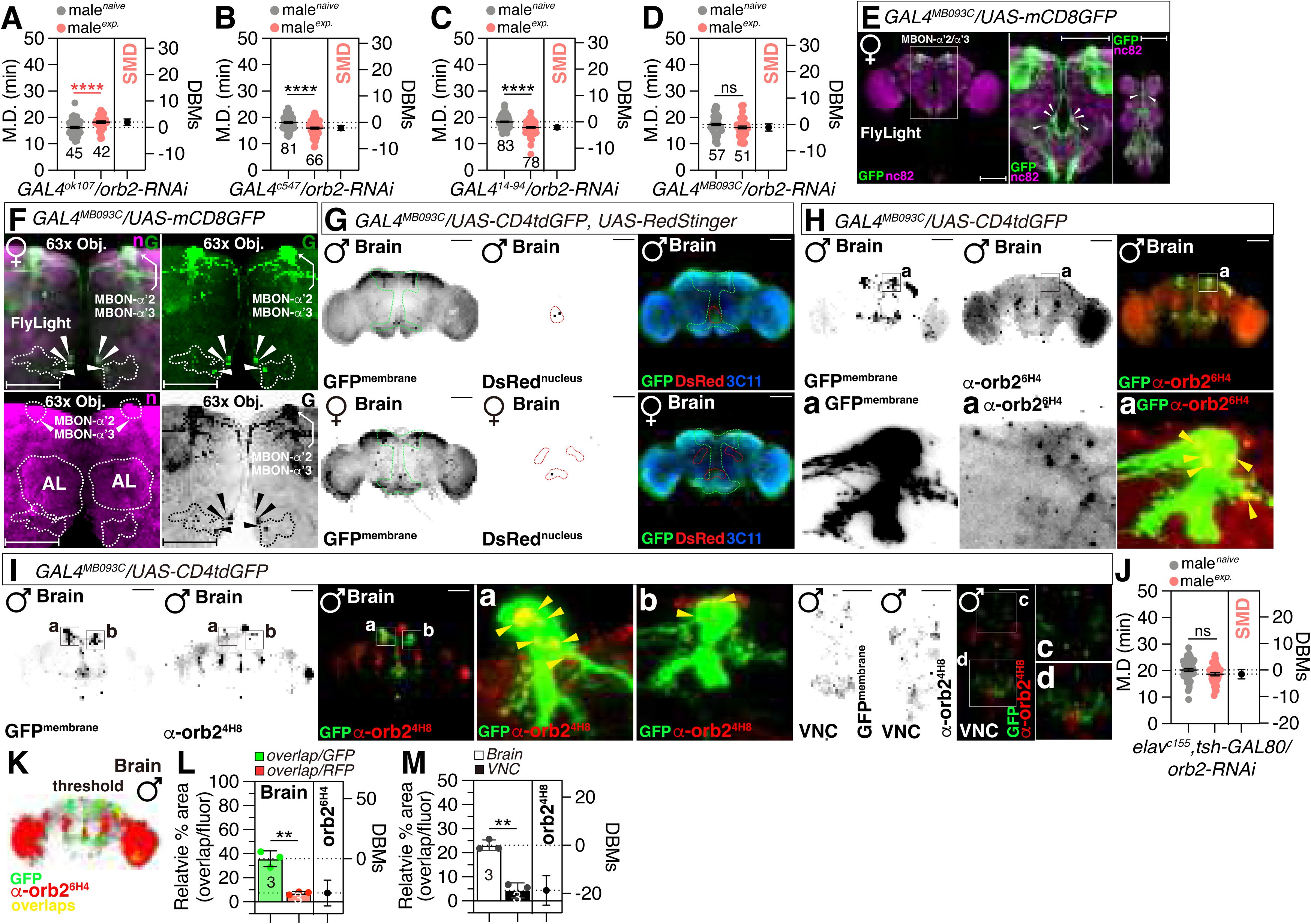
The presence of the Orb2 protein in *GAL4^MB093C+^* cells is necessary for the generation of SELTM. (A-D) SMD assays for *GAL4^ok107^* (A)*, GAL4^c547^* (C)and *GAL4^14-94^* (C) *and GAL4^MB093C^* (D) mediated knockdown of *orb2 via orb2-RNAi*. (E-F) Brain and VNC of male flies expressing *GAL4^MB093C^* with *UAS-CD4tdGFP,* were immunostained with anti-GFP (green) and nc82 (magenta) antibodies. Scale bars represent 100 μm. Boxes indicate the magnified regions to clearly show the expression patterns of neurons in brain labeled by *GAL4^MB093C^* driver (E). And the zoom in the MB-body of interesting patterns (F), utilize white dashed lines to delineate regions of significance and provide explanatory notes using text. The data from Flylight (https://www.janelia.org/project-team/flylight). (G) Brain of male flies expressing *GAL4^MB093C^* together with *UAS-CD4tdGFP, UAS-Redstinger* were immunostained with anti-GFP (green), anti-DsRed (red), and 3c11(blue) antibodies. Scale bars represent 100 μm. Green and red lines indicate the magnified regions of interest presented in the left two panels are presented as a grey scale to clearly show the expression patterns of neurons in brain labeled by *GAL4^MB093C^* driver. For detailed methods, see the METHODS for a detailed description of the immunostaining procedure used in this study. (H) Brain of male flies expressing *GAL4^MB093C^* with *UAS-CD4tdGFP* were immunostained with anti-GFP (green) and anti-orb2^6H4^ (red). Boxes indicate the magnified regions of interest presented in the left two panels are presented as a grey scale to clearly show the expression patterns of neurons in brain labeled by *GAL4^MB093C^* driver, The yellow arrow indicates the precise position of the Orb2 protein within the synapse alongside *GAL4^MB093C+^* cells. (I) Brain and VNC of male flies expressing *GAL4^MB093C^* with *UAS-CD4tdGFP* were immunostained with anti-GFP (green) and anti-orb2^4H8^ (red). Boxes indicate the magnified regions of interest presented in the left two panels are presented as a grey scale to clearly show the expression patterns of neurons in brain labeled by *GAL4^MB093C^*driver. (J) SMD assays for *elav^c155^*mediated knockdown of Orb2 *via orb2-RNAi* together with *tsh-GAL80*. (K-L) Colocalization analysis of GFP and DsRed staining, normalized to total GFP and DsRed areas. Bars represent the mean GFP (green column) and DsRed (red column) fluorescence level with error bars representing SEM. Asterisks represent significant differences, as revealed by the Student’s *t* test and ns represents non-significant difference (*\*p<0.05, **p<0.01, ***p< 0.001, ****p< 0.0001*). The same symbols for statistical significance are used in all other figures. See the METHODS for a detailed description of the colocalization analysis used in this study. (M) Colocalization analysis of DsRed (Orb2) in Brain and VNC, normalized to total DsRed areas, Bars represent the mean Brain (white column) and DsRed (black column) fluorescence level with error bars representing SEM. Asterisks represent significant differences, as revealed by the Student’s *t* test and ns represents non-significant difference (*\*p<0.05, **p<0.01, ***p< 0.001, ****p< 0.0001*). The same symbols for statistical significance are used in all other figures. See the METHODS for a detailed description of the colocalization analysis used in this study.

**Table 1.**
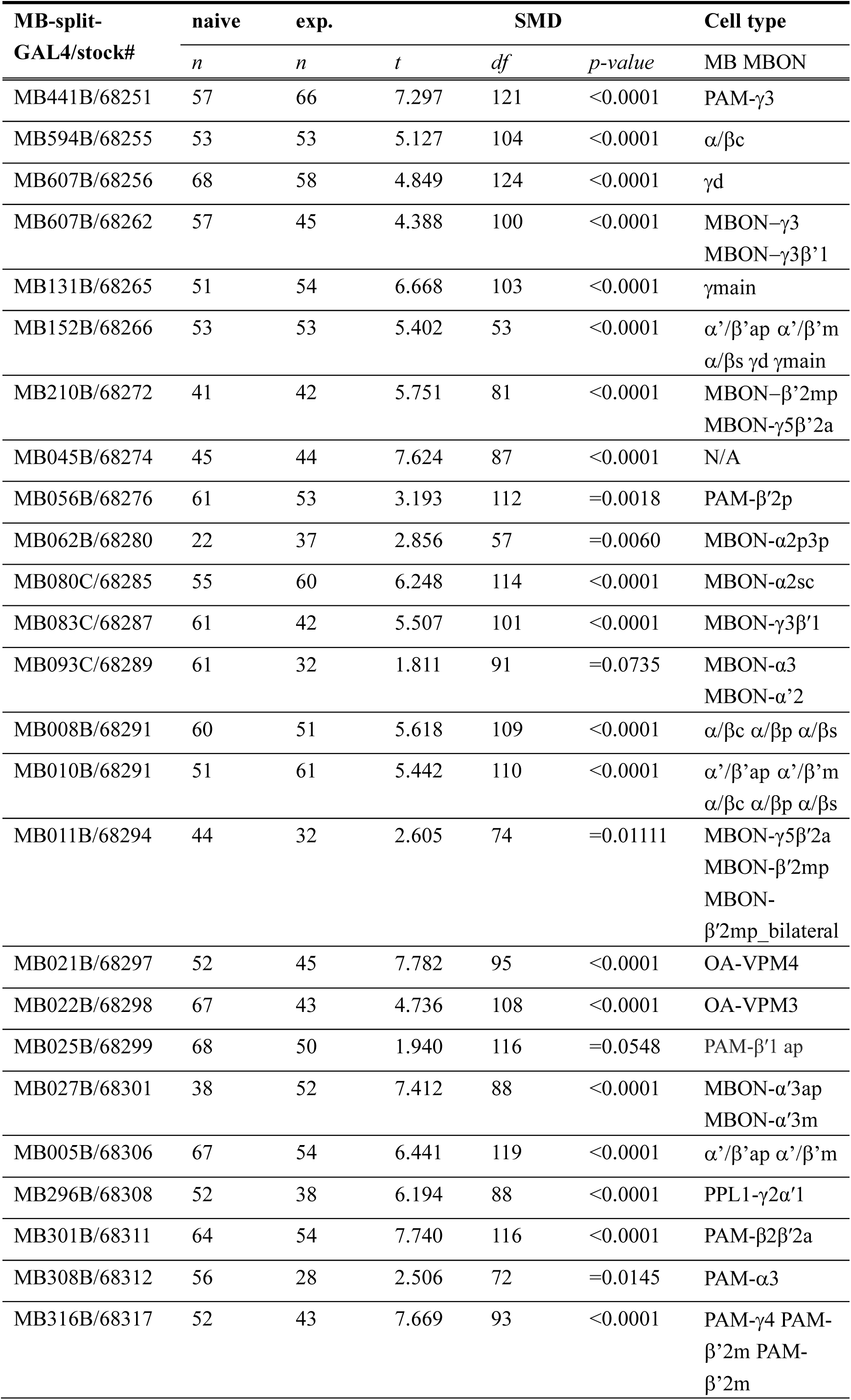
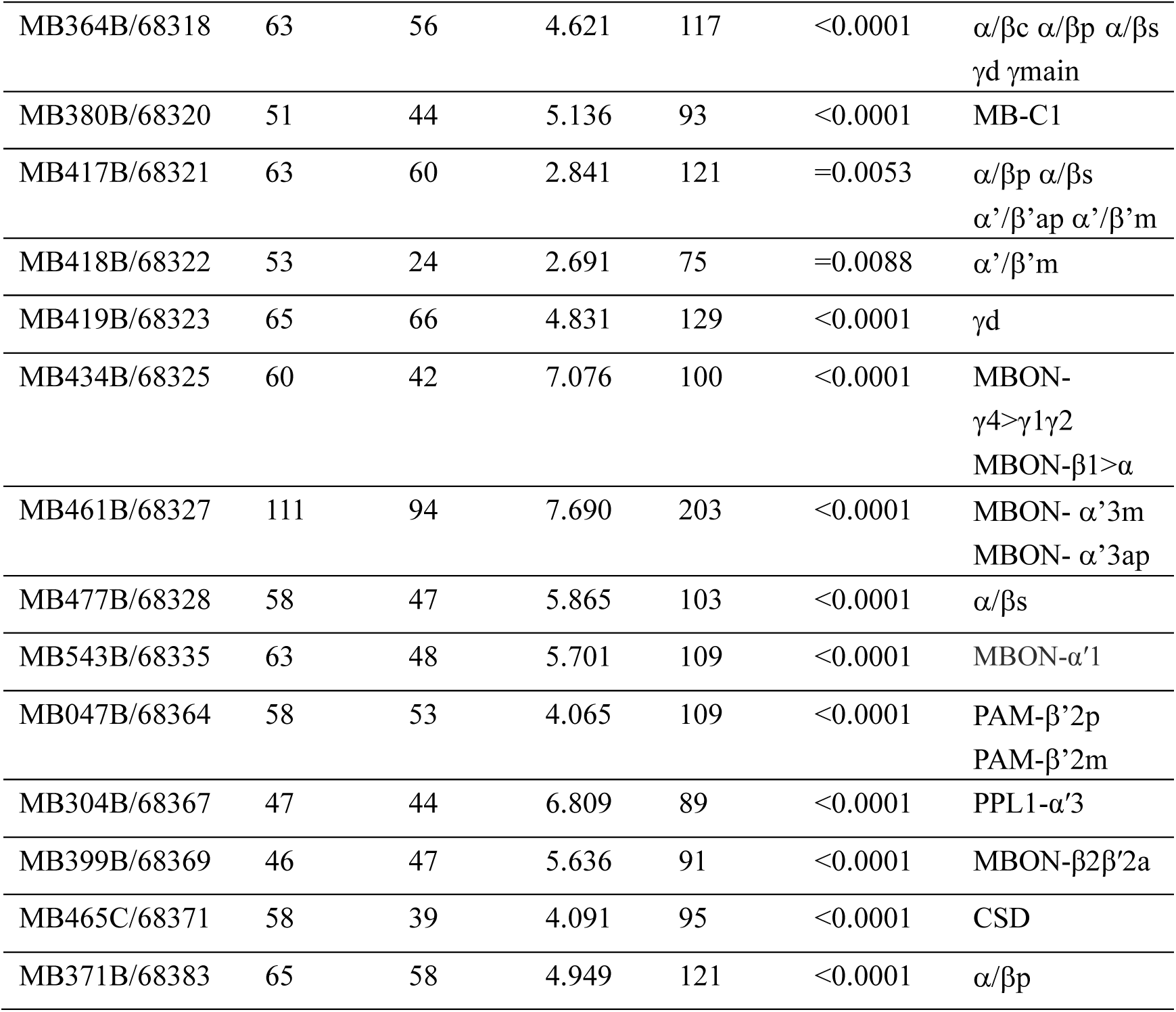
Summary screening of SMD assays for *MB-Split-GAL4* mediated knockdown of Orb2 via *orb2-RNAi* data.

The FlyLight GAL4 expression pattern indicates that *GAL4^MB093C^* exclusively labels three pairs of neurons with cell bodies situated below the AL and above the subesophageal ganglion (SOG) (Fig. 2E)^48^. Each of these neurons extends a vertical fiber to the dorsal brain region, where they branch into dense arborizations within the α lobes of the MB and send sparse terminals to the anterior, middle, and posterior superiormedial protocerebrae in female brains^49^ (Fig. 2F). Our further analysis confirmed the presence of only three pairs of nuclei near the SOG in male brains, whereas female brains exhibit a greater number of nuclei near the AL (Fig. 2G), suggesting subtle sexual dimorphisms in *GAL4^MB093C^*-expressing neurons.

Orb2 expression in *GAL4^MB093C^* neurons was validated by three independent anti-Orb2 antibodies (Fig. 2H-I and Fig. S2D). Remarkably, Orb2 protein forms highly oligomerized puncta in the MB α-lobe area to which *GAL4^MB093C^* neurons project (yellow arrows in Fig. 2H-I and Fig. S2D). We observed leaky expression of *GAL4^MB093C^* in the VNC (panel VNC in Fig. 2I and Fig. S2D), but knockdown of *Orb2* exclusively in the brain still disrupted SMD behavior (Fig. 2J). Additionally, Orb2 puncta did not colocalize with *GAL4^MB093C^*-positive VNC neurons (Fig. 2K-M and Fig. S2E). Notably, the MB α-lobe displays increased colocalization with the Orb2 protein following male sexual experience (Fig. S2F-J), implying that Orb2 accumulation in the MB α-lobe post-experience may enhance memory formation.

The collective results demonstrate that the expression of the Orb2 gene in the three pairs of MB projection neurons is essential for the formation of SELTM with SMD behavior, which we have designated as ’SELTM’. We have designated these three pairs of neurons as ‘Yuelao (YL)’ neurons, a nomenclature derived from Chinese mythology, where ‘Yuelao’ is the deity associated with marriage and love.

### RNA-binding proteins and the CaM-Kinase signaling cascade are indispensable constituents of SELTM

RNA-binding proteins, including Staufen (Stau), Fragile X messenger ribonucleoprotein 1 (Fmr1), and Pumilio (Pum), are integral to the molecular underpinnings of LTM in *Drosophila*^50–53^ (Fig. 3A). Specifically, Fmr1 expression is a prerequisite for SELTM in YL neurons (Fig. 3B-D, S3A-B). Colocalization analysis via Fly SCope suggests that Orb2 co-expresses with Stau, Pum, and Fmr1 in the Kenyon cell population (Fig. 3E-G), indicating that YL neurons may employ distinct molecular mechanisms for SELTM compared to those associated with other sensory modalities.

**Fig 3.**
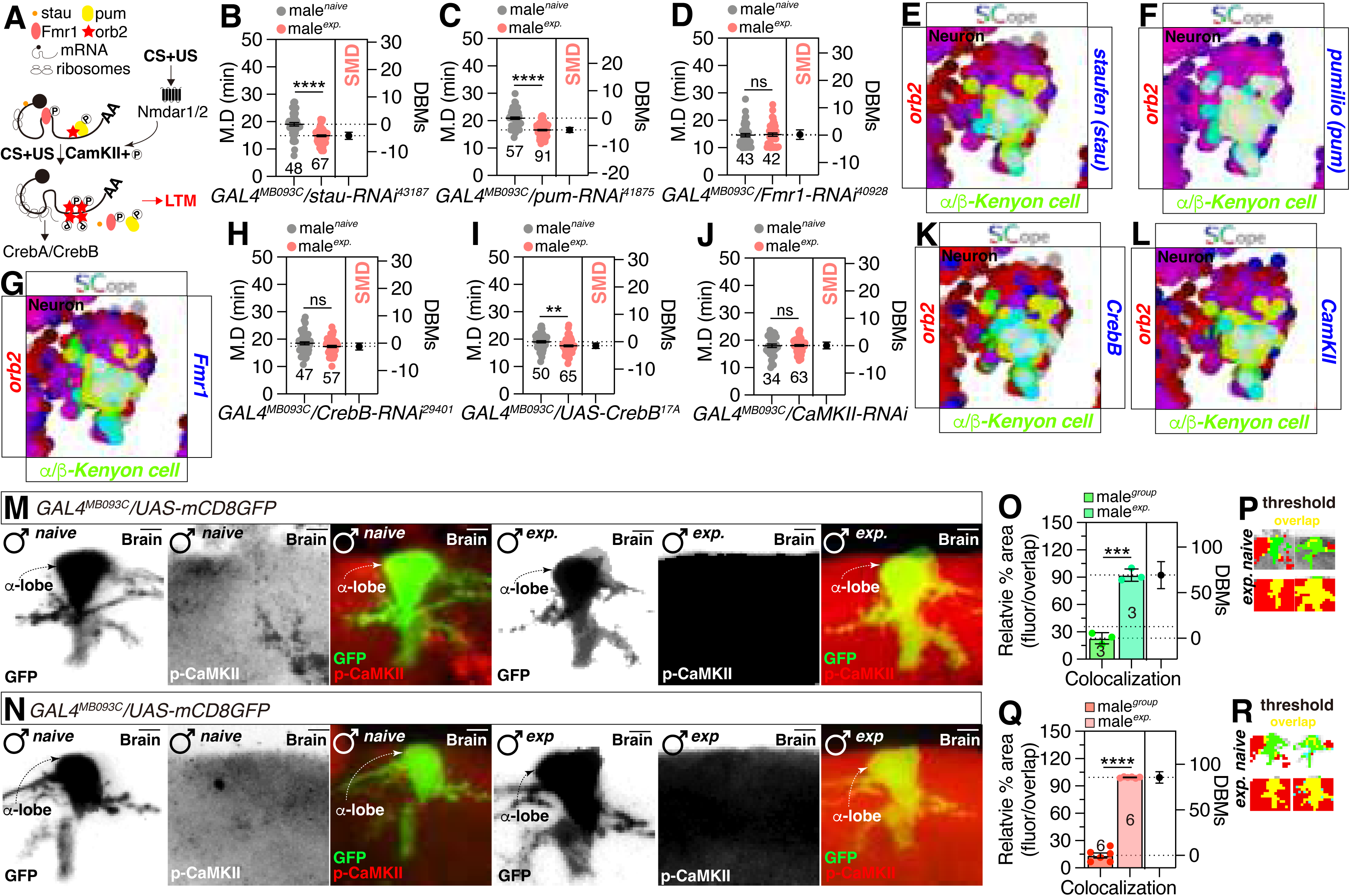
The presence of the CPEB proteins in YL neuron is necessary for the manifestation of SELTM. (A) Illustration depicting the interaction CPEB and mRNA Creating and strengthening LTM related to SMD under conditioned stimulus (CS) and unconditioned stimulus (US) circumstances, The orange dot represents the stau, the yellow dot represents the of pum. Fmr1 is represented by the pink oval, while Orb2 is represented by the red star. (B-D) SMD assays for *GAL4^MB093C^* mediated knockdown of stau, pum and Fmr1 *via stau-RNAi^i43187^* (B) *pum -RNAi^41875^* (C) *and Fmr1 -RNAi^40928^* (D). (E-G) Single-cell RNA sequencing (SCOPE scRNA-seq) datasets reveal cell clusters colored by expression of *orb2* (red) *stau* (blue) and α/β-Kenyon cell (green) in mushroom body neurons (E), expression of *orb2* (red) *pum* (blue) and α/β-Kenyon cell (green) in mushroom body neurons (F) and expression of *orb2* (red) Fmr1 (blue) and α/β-Kenyon cell (green) in mushroom body neurons (G). (H) SMD assays for *GAL4^MB093C^* mediated knockdown of CrebB, *via CrebB-RNAi^29401^*. (I) SMD assays for *GAL4^MB093C^* overexpress of CrebB via *UAS-CrebB^17A^*. (J) SMD assays for *GAL4^MB093C^* mediated knockdown of CaMKII via *CaMKII-RNAi.* (K-L) Single-cell RNA sequencing (SCOPE scRNA-seq) datasets reveal cell clusters colored by expression of *orb2* (red) *CrebB* (blue) and α/β-Kenyon cell (green) in mushroom body neurons (K) and expression of *orb2* (red) CaMKII (blue) and α/β-Kenyon cell (green) in neurons (L). (M-N) MB-α/β-lobe of male flies expressing *GAL4^MB093C^* with *UAS-CD4tdGFP* were immunostained with anti-GFP (green) and anti-orb2^6H4^ (red). (O-R) Colocalization analysis of GFP and DsRed staining, normalized to total GFP and DsRed areas. Bars represent the mean GFP, green column (O) and DsRed, red column (Q). To effectively illustrate the phenomenon of colocalization, the outcomes of GFP (P) and RFP (R) are presented on the right side, while the terms “naive” and “exp.” are noted in the picture. See the METHODS for a detailed description of the colocalization analysis used in this study.

Cyclic AMP response element-binding protein (CREB)-responsive transcription has been established as a pivotal mechanism in LTM formation^54,55^. In *Drosophila*, CrebA and CrebB encode CREB family proteins, and their knockdown of CrebB in YL neurons disrupts SELTM (Fig. 3H). However, overexpression of CrebB does not influence SELTM (Fig. 3I), suggesting that the presence of CrebB is more critical than its expression level for SELTM generation. *Drosophila* Calcium/Calmodulin-dependent Kinase II (CaMKII) and Calcium/Calmodulin-dependent Kinase I (CaMKI), a homolog of the mammalian αCaMKII and CaMKI, is synthesized at synapses and its mRNA translation is regulated by cytoplasmic polyadenylation element–binding (CPEB) proteins at synapses, a process crucial for LTM formation^56–59^. Knockdown of CaMKI or CaMKII disrupts SELTM (Fig. 3J and. Fig. S3C). The elevated expression of the wild-type CaMKII protein does not exert a significant influence on SELTM (Fig. S3D). Conversely, the introduction of a mutant variant of CaMKII leads to the impairment of SELTM (Fig. S3E-F).

It is worth mentioning that knockdown of both CaMKII and CaMKI disrupts LTM for SMD in YL neurons (Fig. 3J and Fig. S3C) and the Fly SCope data indicate that all LTM-related genes are highly enriched in MB Kenyon cells co-expressing Orb2 (Fig. 3K-L), suggesting that the co-expression of these genes with Orb2 does fully account for the specificity of SELTM formation (Fig. S3G-L for genetic control experiments).

### The signaling pathway mediated by Nmadr1 is crucial for the SELTM formation

In *Drosophila melanogaster*, the NMDAR1 (N-methyl-D-aspartate receptor1) and its homolog, NMDAR2, are pivotal components of the glutamatergic signaling system, which is integral to various brain functions, including memory formation. NMDARs are ionotropic glutamate receptors that mediate rapid excitatory synaptic transmission and are implicated in long-term potentiation (LTP), a form of synaptic plasticity that is considered a molecular correlate of learning and memory^60,61^. The NMDARs-CamKI/II signaling cascade is a critical component of LTM (Fig. 4A)^62^. RNA interference-mediated knockdown of *Nmdar2*, but not *Nmdar1*, disrupts SELTM (Fig. 4B-C), despite both receptors co-expressing in Orb2-positive MB Kenyon cells (Fig. 4D- E). Notably, RNAi knockdown of NMDAR2 in *GAL4^ok1^*^07^*-*positive MB neurons does not affect SELTM (Fig. 4F), suggesting that NMDAR2 expression in YL neurons, but not in other MB neurons, is required for the generation of SELTM (Fig. S4A-B for genetic control experiments).

**Fig 4.**
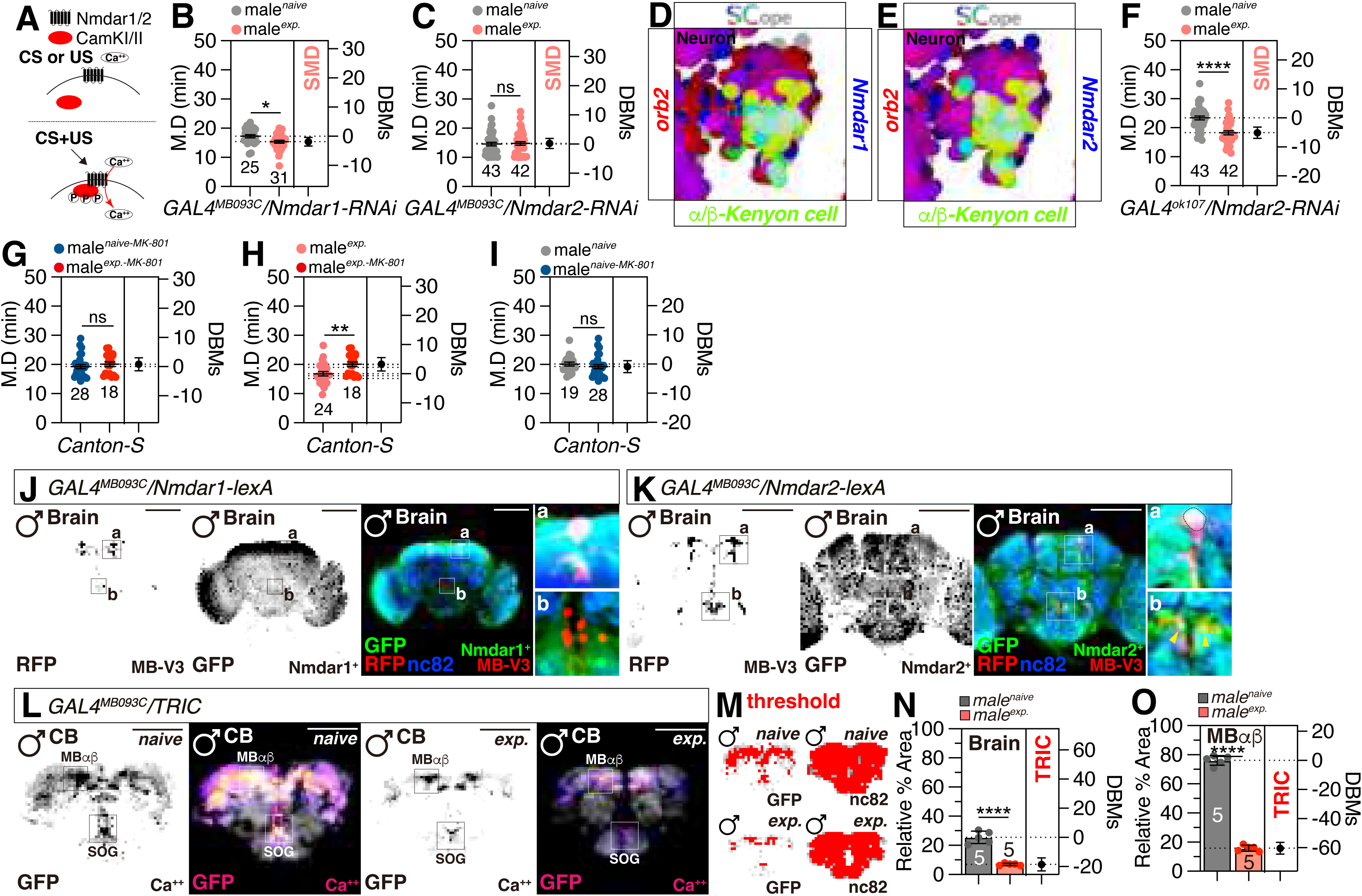
The presence of the NMDAR2 in YL neuron is necessary for the manifestation of SELTM. (A) Illustration depicting the interaction between NMDARs and CaMKI/II under conditioned stimulus (CS) and unconditioned stimulus (US) circumstances. (B-C) SMD assays for *GAL4^MB093C^* mediated knockdown of NMDAR1 and NMDAR2 via *Nmdar1-RNAi* (B) and *Nmdar2-RNAi* (C). (D-E) Single-cell RNA sequencing (SCOPE scRNA-seq) datasets reveal cell clusters colored by expression of *orb2* (red) *Nmdar1* (blue) and α/β-Kenyon cell (green) in mushroom body neurons (D) and expression of *orb2* (red) *Nmdar2* (blue) and α/β-Kenyon cell (green) in mushroom body neurons (E). (F) SMD assays for *GAL4^ok107^* mediated knockdown of NMDAR2 via *Nmdar2-RNAi*. (G-I) SMD assays for *Canton-s* with MK-801 feeding. (J-K) Brain of male flies expressing *Nmdar1-LexA* (J) or *Nmdar2-LexA* (K) and *GAL4^MB093C^* together with *CD4tdGFP-LexAop, UAS-Redstinger* were immunostained with anti-GFP (green), anti-DsRed (red), and nc82 (blue) antibodies. Scale bars represent 100 μm. Boxes indicate the magnified regions of interest presented in the left two panels are presented as a grey scale to clearly show the expression patterns of neurons in brain labeled by *GAL4^MB093C^* driver and the right two panels are zoom in the boxes indicate the magnified regions to clearly shows the overlapping. For detailed methods, see the METHODS for a detailed description of the immunostaining procedure used in this study. (L) Different levels of neural activity of the brain as revealed by the TRIC system in naïve and experienced flies. Male flies expressing *GAL4^MB093C^* along with *UAS-UAS-IVS-mCD8RFP, LexAop2-mCD8GFP, nSyb-MKII-LexA, UAS-p65AD::CaM* and were dissected after 5 days of growth (mated male flies had 1-day of sexual experience with virgin females). The dissected brains were then immunostained with anti-GFP (fire), anti-nc82 (grey). GFP is pseudo-colored as “red hot”. Boxes indicate the magnified regions of interest presented in the left panels. Scale bars represent 100 μm. And zoom in the boxes indicate the magnified regions, MB-α/β-lobe (L). For detailed methods, see the METHODS for a detailed description of the immunostaining procedure used in this study. (M-O) Quantification was performed for the DsRed and GFP fluorescence in CB, Mainly SOG (K) and MB-α/β-lobes (L) between naïve and experienced male flies. The GFP fluorescence was normalized relative to the fluorescence of thenc82. The conditions of flies are described above: naïve, naïve male flies; exp., male flies with sexual experience. Bars represent the mean of the normalized GFP fluorescence level with error bars representing the SEM. Asterisks represent significant differences, as revealed by the Student’s *t* test and ns represents non-significant difference (*\*p < 0.05, **p < 0.01, ***p < 0.001*). See the METHODS for a detailed description of the fluorescence intensity analysis used in this study.

Dizocilpine, also known as MK-801, is a selective antagonist of N-methyl-D-aspartate receptors (NMDARs), which are a subclass of ionotropic glutamate receptors critical to the functioning of the CNS ^63^. MK-801 exerts its antagonistic effect by binding to two distinct sites within the NMDAR-ion channel complex, thereby attenuating glutamatergic neurotransmission through NMDARs. The application of MK-801 to block NMDAR activity results in the disruption of SELTM (Fig. 4G). Our subsequent analysis, conducted under both naive and experienced conditions with and without MK-801 (0.3 mM) administration, revealed that MK-801 specifically suppresses the reduction in MD induced by sexual experience (Fig. 4H and 4I). Collectively, these findings indicate that the specific inhibition of NMDARs by MK-801 interferes with the consolidation of SELTM in subjects with prior sexual experience.

Genetic intersection analyses corroborated that Nmdar1 is not expressed with YL neurons (Fig. 4J), whereas Nmdar2 is expressed in a single pair of YL neurons in both male and female brains (indicated by yellow arrows in Fig. 4K and Fig. S4C). These data suggests that the NMDAR2-CaM-Kinases cascade in these specific YL neurons is a pivotal component for the generation of SELTM. CaMKII phosphorylation of various synaptic proteins is pivotal in modulating their structural and functional dynamics, which are pivotal for synaptic plasticity^62^. Intriguingly, the overall calcium levels within YL neurons were markedly reduced following sexual experience in males (Fig. 4L-O and Fig. S4D-H). The observed reduction in calcium levels within YL neurons suggests that the inhibitory function of the neuronal signaling pathway plays a crucial role in the formation of SELTM.

### Short neuropeptide F (sNPF) inputs to YL neurons connect with NMDAR2 inputs to release glutamate for LTM formation

Previously, we elucidated that sNPF-sNPF-R signaling within the AL and the MB dorsal accessory calyx (dAC) is pivotal for modulating SMD behavior ^64^. The male-specific sNPF-sNPF-R signaling circuits operate as an accumulator, processing information related to sexual experiences. The integration of this information into memory circuits is essential for interval timing, with accumulator circuits being proposed to be interconnected with memory circuits^39–41,43,65,66^. It is hypothesized that sNPF-R may serve as the neural interface that bridges the accumulator circuits into memory circuits (Fig. 5A).

**Fig 5.**
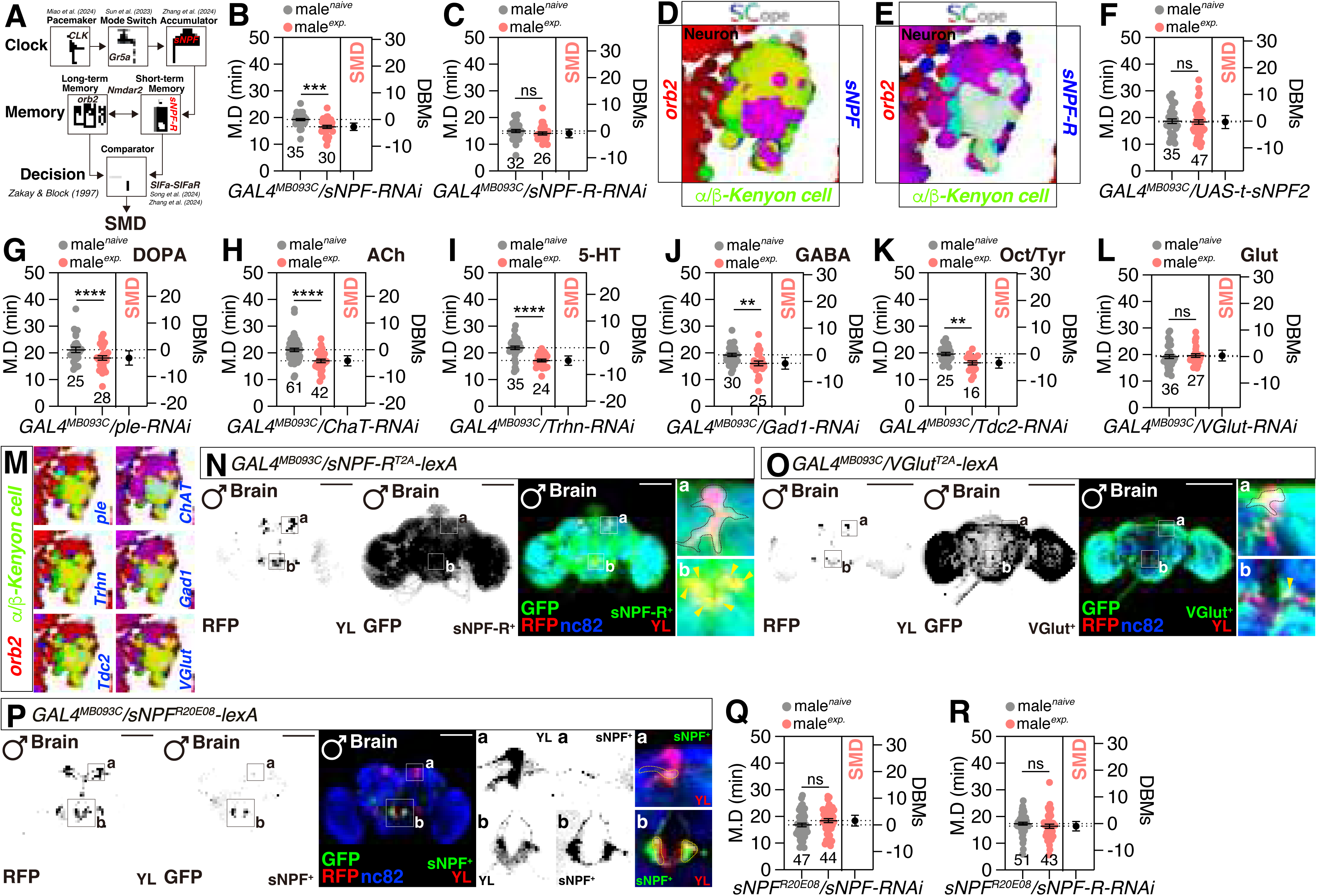
YL neurons express sNPF-R and function as glutamatergic neurons. (A) The behavior of SMD is determined by the behavioral hypothesis model put out by Interval timing, wherein all model components are abstracted to enhance reader comprehension. CLK neurons function as pacemakers, Gr5a cells act as switches, and sNPF neurons serve as accumulators. The focus of this article is the memory component, which consolidates memories and transmits it to the downstream decision-making neurons known as SIFa neurons. (B-C) SMD assays for *GAL4^MB093C^* mediated knockdown of sNPF and sNPF-R via*sNPF-RNAi* (B) and *sNPF-R-RNAi* (C). (D-E) Single-cell RNA sequencing (SCOPE scRNA-seq) datasets reveal cell clusters colored by expression of orb2 (red) sNPF (blue) and α/β-Kenyon cell (green) in mushroom body neurons (D) and *orb2* (red) *sNPF-R* (blue) and α/β-Kenyon cell (green) in mushroom body neurons (E). (F) SMD assays for *GAL4^MB093C^* expression for self-coupling of t-sNPF-2 via *UAS-t-sNPF-2-SEC*. (G-L) SMD assays for *GAL4^MB093C^* mediated knockdown of *ple* (G), ChaT (H), Trhn (I), Gad1 (J), Tdc2 (K) and VGlut (L) via *Neurotransmitters -RNAi.* (M) Single-cell RNA sequencing (SCOPE scRNA-seq) datasets reveal cell clusters colored by expression of orb2 (red) *ple, ChaT, Thrn, Gad1, Tdc2* and *VGlut* (blue) and α/β-Kenyon cell (green) in neurons (mushroom body). (N-P) Brain of male flies expressing *sNPF-R^T2A^-LexA* (N), *VGlut^T2A^-LexA* (O) or *sNPF^20E06^-LexA* (P) and *GAL4^MB093C^* together with *CD4tdGFP-LexAop, UAS-Redstinger* were immunostained with anti-GFP (green), anti-DsRed (red), and nc82 (blue) antibodies. Scale bars represent 100 μm. Boxes indicate the magnified regions of interest presented in the left two panels are presented as a grey scale to clearly show the expression patterns of neurons in brain labeled by *GAL4^MB093C^* driver and the right two panels are zoom in the boxes indicate the magnified regions to clearly shows the overlapping. For detailed methods, see the METHODS for a detailed description of the immunostaining procedure used in this study. (Q-R) SMD assays for *sNPF^20E06^-GAL4* mediated knockdown of sNPF and sNPF-R via *sNPF-RNAi* (Q) and *sNPF-R-RNAi* (R).

Knockdown of sNPF-R, but not sNPF, within YL neurons disrupts SELTM (Fig. 5B-C), and sNPF-R is exclusively expressed with Orb2 in MB Kenyon cells (Fig. 5D-E). Furthermore, the overexpression of membrane-tethered sNPF in YL neurons disrupts SELTM (Fig. 5F and Fig. S5A-E), indicating that the proximity of sNPF to YL neurons is a critical determinant for SELTM formation (Fig. S5F for genetic control experiments).

Although sNPF has been demonstrated to act as a neuromodulator within the MB for olfactory memory^67^, the specific neurotransmitters (NTs) that synergize with sNPF in MB memory circuits remain elusive. Utilizing RNA interference-mediated knockdown of essential enzymes involved in NT synthesis, we identified that glutamate release from YL neurons is indispensable for LTM induction (Fig. 5G-M and Fig. S5G-L for genetic control experiments). Additionally, YL neurons do not appear to receive inputs from serotonergic neurons through 5-HT receptors, despite the established requirement of serotonin receptors in MB for learning and memory formation^68–71^ (Fig. S5M-P).

Recently generated knockin *sNPF-R^T2A^-lexA* driver expresses in all three pairs of YL neurons^72^ (Fig. 5N), whereas *Vglut^T2A^-lexA* is expressed in only one pair of medial neurons (Fig. 5O). The localization of VGlut-positive YL neurons corresponds to that of Nmdar2-positive YL neurons (Fig. 4K), suggesting that this specific YL neuron pair receives input signals through NMDAR2 and sNPF-R, subsequently releasing glutamate into the MB α/β lobes (Fig. S5Q).

Neuropeptide sNPF is known to be broadly expressed throughout the brain and is involved in local neuromodulatory functions^73^. To identify the local sNPF-expressing neurons that interact with YL neurons, we screened GAL4 strains that drive sNPF expression and identified the *sNPF^R20E^*^08^ driver, which is adjacent to YL neurons (Fig. 5P). These *sNPF^R20E^*^08^ neurons partially overlaps with YL neurons and also locate closely with YL neurons in the vicinity of the SOG and the MB α/β lobes. We designated this group of sNPF-expressing local neurons as “Weigu (WG)” neurons, inspired by the Chinese legend of a man who encountered Yue Lao, the deity of marriage. Knockdown of sNPF and sNPF-R in WG neurons abolishes SELTM (Fig. 5Q-R), indicating that WG neurons co-express sNPF and sNPF-R near YL neurons.

These findings collectively suggest that local sNPF inputs from WG neurons to sNPF-R-expressing YL neurons, in conjunction with NMDAR2, are essential for the generation of SELTM behavior (Fig. S5Q).

### The rise in synaptic plasticity of YL neurons is accompanied by a reduction in intracellular calcium concentrations

The NMDAR-CaMKII cascade is well-established as a critical regulator of synaptic plasticity^6^^2,74,75^. To investigate whether the NMDAR2-CaM-Kinases signaling pathway in YL neurons (Movie 1) undergoes synaptic plasticity during memory formation, we utilized the DenMark and syt.eGFP markers to assess the morphology of dendrites and presynaptic terminals^76^. Our findings revealed a substantial augmentation in the area occupied by YL neurons’ dendrites and presynaptic terminals throughout the brain (Fig. 6A and S6A-C), with a more pronounced increase observed in the MB and SOG regions (Fig. 6B-G).

**Fig 6.**
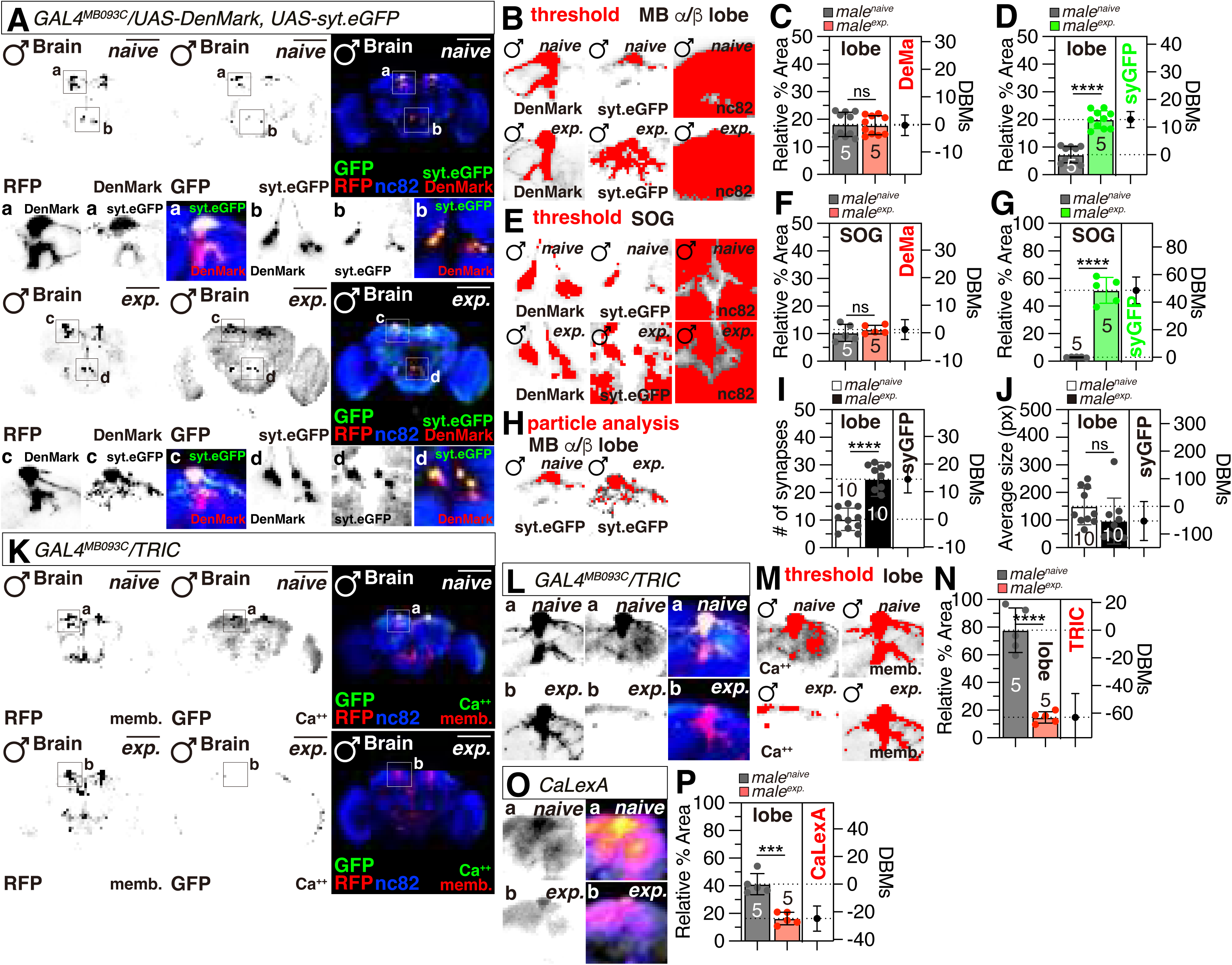
Sexual experiences induce widespread alterations in calcium dynamics subsequent to synaptic plasticity within YL neurons. (A) Male flies brain expressing *GAL4^MB093C^* drivers together with *UAS-Denmark* and *UAS-syteGFP* were immunostained with anti-GFP (green), anti-DsRed (red) and anti-nc82 (blue) antibodies. Scale bars represent 100 μm. Boxes indicate the magnified regions of interest presented in the left and bottom panels. Dashed circles indicate the neurons where *GAL4^MB093C^* form interconnected networks. The left two panels are presented as a grey scale to clearly show the dendritic and synaptic terminal of neurons labeled by *GAL4^MB093C^* driver. (B-G) Quantification of relative value for synaptic area, which is formed by *GAL4^MB093C^* in MB α/β-lobe (B-D) and SOG (E-G), between naïve and experienced male flies. The possible synaptic interactions in naïve and experienced male flies. The small panels are presented as a red scale to show the GFP and DsRed fluorescence marked by threshold function of ImageJ. And quantification was performed for the GFP and DsRed fluorescence in these regions between naïve and experienced male flies. The GFP and DsRed fluorescence was normalized relative to the fluorescence of the nc82. The conditions of flies are described above: naïve, naïve male flies; exp., male flies with sexual experience. Bars represent the mean of the normalized GFP fluorescence level with error bars representing the SEM. Asterisks represent significant differences, as revealed by the Student’s *t* test and ns represents non-significant difference (*\*p < 0.05, **p < 0.01, ***p < 0.001*). See the METHODS for a detailed description of the fluorescence intensity analysis used in this study. (H-J) Quantification of relative value for synaptic particle, which is formed by *GAL4^MB093C^* in MB α/β lobe, between naïve and experienced male flies. The possible synaptic particle interactions in naïve and experienced male flies. The small panels are presented as a red scale to show the GFP fluorescence marked by threshold function of ImageJ (H). And quantification was performed for the GFP particle number (I) and average size (J) in these regions between naïve and experienced male flies. Asterisks represent significant differences, as revealed by the Student’s *t* test and ns represents non-significant difference (*\*p < 0.05, **p < 0.01, ***p < 0.001*). See the METHODS for a detailed description of the fluorescence intensity analysis used in this study. (K-L) Different levels of neural activity of the brain as revealed by the TRIC system in naïve and experienced flies (K). Male flies expressing *GAL4^MB093C^* along with *UAS-UAS-IVS-mCD8RFP, LexAop2-mCD8GFP, nSyb-MKII-LexA, UAS-p65AD::CaM* and were dissected after 5 days of growth (mated male flies had 1-day of sexual experience with virgin females). The dissected brains were then immunostained with anti-GFP (green), anti-RFP(red) and anti-nc82 (blue). GFP is pseudo-colored as “red hot”. Boxes indicate the magnified regions of interest presented in the left panels. The left three panels are presented as a grey scale to clearly show the expression patterns of neurons in brain labeled by *GAL4^MB093C^* driver. Scale bars represent 100 μm. And zoom in the boxes indicate the magnified regions, MB-α/β-lobe (L). For detailed methods, see the METHODS for a detailed description of the immunostaining procedure used in this study. (M-N) Quantification was performed for the DsRed and GFP fluorescence in MB-α/β-lobe between naïve and experienced male flies. The GFP fluorescence was normalized relative to the fluorescence of the DsRed. The conditions of flies are described above: naïve, naïve male flies; exp., male flies with sexual experience. Bars represent the mean of the normalized GFP fluorescence level with error bars representing the SEM. Asterisks represent significant differences, as revealed by the Student’s *t* test and ns represents non-significant difference (*\*p < 0.05, **p < 0.01, ***p < 0.001*). See the METHODS for a detailed description of the fluorescence intensity analysis used in this study. (O) Different levels of neural activity of the MB-α/β-lobe as revealed by the CaLexA system in naïve, single and experienced flies. Male flies expressing *GAL4^MB093C^* along with *LexAop-CD2-GFP, UAS-mLexA-VP16-NFAT and LexAop-CD8-GFP-A2-CD8-GFP* were dissected after 5 days of growth (mated male flies had 1-day of sexual experience with virgin females). The dissected brains were then immunostained with anti-GFP (green) and anti-nc82 (blue). GFP is pseudo-colored as “red hot”. Boxes indicate the magnified regions of interest presented in the bottom panels. Scale bars represent 100 μm. (P) Quantification was performed for the GFP fluorescence in MB-α/β-lobe between naïve and experienced male flies. The GFP fluorescence was normalized relative to the fluorescence of the nc82.The conditions of flies are described above: naïve, naïve male flies; exp., male flies with sexual experience. Bars represent the mean of the normalized GFP fluorescence level with error bars representing the SEM. Asterisks represent significant differences, as revealed by the Student’s *t* test and ns represents non-significant difference (*\*p < 0.05, **p < 0.01, ***p < 0.001*). See the METHODS for a detailed description of the fluorescence intensity analysis used in this study.

A detailed particle analysis within the MB region demonstrated a marked increase in the number of synapses proximate to the α/β lobes of the MB, while the average size of these synapses remained unchanged (Fig. 6H-J). These data indicate that the increase of CaMKII phosphorylation within YL neurons induced by sexual experiences enhances their synaptic plasticity, thereby facilitating the establishment of SELTM.

To further corroborate the calcium dynamics during synaptic plasticity, we utilized a transcriptional reporter of intracellular calcium (TRIC)^77^. Our results revealed a significant decrease in the calcium levels of YL neurons following the onset of synaptic plasticity (Fig. 6K-N and Fig. S6D-E), particularly in the α/β lobes of the MB (Fig. 6L-N). Additionally, the intracellular calcium reporter CaLexA displayed concordant results^78^ (Fig. 6O-P and S6F-H), suggesting that sexual experience indeed leads to a reduction in intracellular calcium levels within YL neurons. Collectively, these data indicate that the sexual experience of males alters the calcium signaling cascade within YL neurons, which in turn promotes increased synaptic plasticity and SELTM (Fig. S6Q).

### YL neurons are specialized for the generation of SELTM

As previously documented, the generation of SMD behavior in male *Drosophila* is significantly influenced by taste and pheromonal cues, which are detected and processed through Gr5a-positive neurons specialized in sugar sensing^37^. The YL neurons, which are situated near the SOG, which have been identified as a primary gustatory processing center ^79,80^. Consequently, we hypothesize that the YL neurons are specialized in the generation of contact-based taste- and pheromone-specific appetitive LTM.

To validate this hypothesis, we conducted a series of olfactory conditioning experiments using a T-maze apparatus (Fig. 7A)^81^. Untrained flies demonstrated a preference for both odor A and B, with approximately a 50% chance of choice for each (Fig. 7B and Fig. S7A-C). Upon conditioning with an electric shock paired with one odor, the flies developed a strong aversion to that odor, avoiding it for up to 12 hours post-training (Fig. 7C-E). Surprisingly, the olfactory LTM was still observed in flies with targeted knockdown of Orb2 or NMDAR2 in the YL neurons (Fig. 7F-K), suggesting that the function of Orb2 and NMDAR2 in YL neurons is not essential for olfactory LTM.

**Fig 7.**
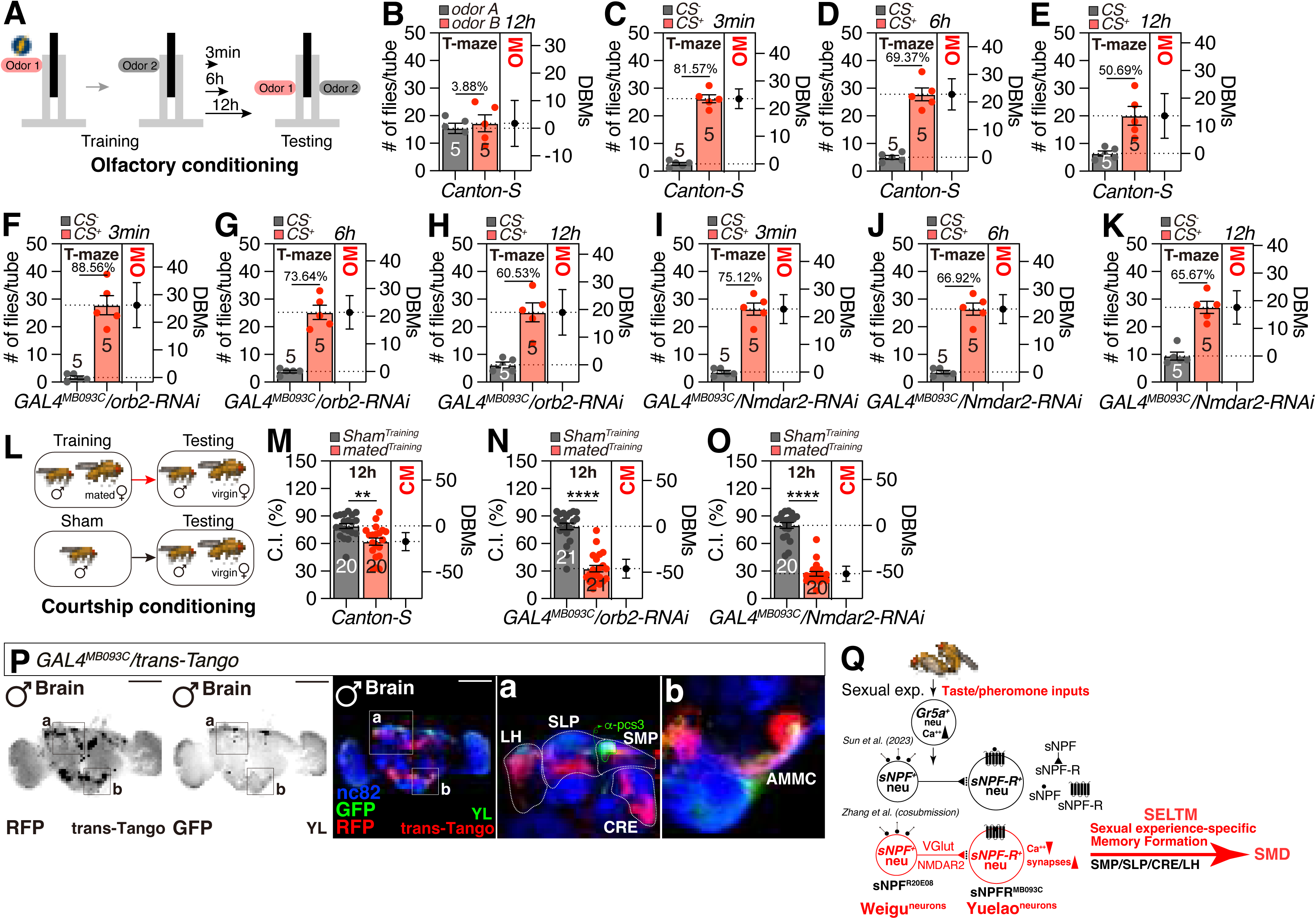
YL neurons do not play a role in the establishment and strengthening of olfactory and courtship memories. |(A) Device for testing olfactory memory During the training session in the T-maze, gas 1 is administered and electrical stimulation is delivered, whereas gas 2 is not administered. Following the completion of training (as shown in the picture), the fruit flies are subjected to several memory tests at various intervals. (B-E) T-maze assays for, the no electrical stimulation *Canton-S* (B), 3 mins interval *Canton-S* (C), 6h interval *Canton-S* (D) and 12h interval *Canton-S* (E). All the interval is after the necessary rest period after training. The numbers on the horizontal line in the figure are MPI, see the METHODS for a detailed description of the fluorescence intensity analysis used in this study. (F-H) T-maze assays for *GAL4^MB093C^* mediated knockdown of orb2 via *orb2-RNAi*., 3 mins interval (F), 6h interval (G) and 12h interval (H). All the interval is after the necessary rest period after training. The numbers on the horizontal line in the figure are MPI, see the METHODS for a detailed description of the fluorescence intensity analysis used in this study. (I-K) T-maze assays for *GAL4^MB093C^* mediated knockdown of NMDAR2 via *Nmdar2-RNAi*., 3 mins interval (I), 6h interval (J) and 12h interval (K). All the interval is after the necessary rest period after training. The numbers on the horizontal line in the figure are MPI, see the METHODS for a detailed description of the fluorescence intensity analysis used in this study. (L) Tarin male flies underwent courtship memory testing by being trained with mated female flies, while sham male flies did not get any training with female flies for a duration of 5 hours. Following a 12-hour period, the male flies underwent memory testing. (M-O) t courtship memory assays for *Canton-S* (M) *GAL4^MB093C^* mediated knockdown of Orb2 via *orb2-RNAi* (N) *GAL4^MB093C^* mediated knockdown of NMDAR2 via *Nmdar2-RNAi* (O). The numbers on the horizontal line in the figure are MPI, see the METHODS for a detailed description of the fluorescence intensity analysis used in this study. (P) Male flies brain expressing *GAL4^MB093C^*drivers with *UAS-myrGFP.QUAS-mtdTomato-3xHA; trans-Tango* were immunostained with anti-GFP (green), anti-DsRed (red) and anti-nc82 (blue) antibodies. Scale bars represent 100 μm. Boxes indicate the magnified regions of interest presented in the left and the two panels on the right display magnified regions. The dashed lines in the zoomed in figure on the right is employed to highlight possible postsynaptic connections. (Q) Illustration depicting the SMD behavior of male Drosophila is dependent on sexual experience. It has been shown that sexual experience is used as taste input, The source of this conclusion is indicated in the figure. taste and pheromonal information from Gr5a neurons transmit information about the male’s sexual experience to the sNPF-sNPF-R circuits. These circuits are then processed by the YL neurons, which WG neurons, being sNPF neurons, receive signals from upstream sources and send information to downstream YL neurons through the use of VGlut and sNPF. YL neurons subsequently transmit taste-specific information. The process of memory formation involves the activation of the superior medial prefrontal cortex (SMP), the subgenual anterior cingulate cortex (SLP), the caudal retrosplenial cortex (CRE), and the lateral hypothalamus (LH), which in turn influences self-monitoring and decision-making behavior.

Additionally, we observed distinct LTM in male flies that were repeatedly rejected by females during courtship, indicating the involvement of multisensory inputs in mediating aversive memory^82–84^ (Fig. 7L-M). Interestingly, knockdown of Orb2 or NMDAR2 in YL neurons still resulted in strong LTM for courtship conditioning up to 12 hours (Fig. 7N-O), indicating that YL neurons are not necessary for the generation of aversive LTM.

To further elucidate the neural circuits connected to YL neurons, we utilized transsynaptic mapping using trans-Tango, a method that labels postsynaptic partners of YL neurons^85^. Our findings revealed that the secondary order YL neurons primarily project to dorsolateral brain regions, including the crepine (CRE), superior medial protocerebrum (SMP), lateral protocerebrum (SLP), and lateral horn (LH)^86^.

Specifically, the LH, which contains local neurons (LHLN), input neurons (LHIN), and output neurons (LHON) involved in memory formation and retrieval, has been recognized as a multimodal ventral zone which is known to relay multisensory inputs from the visual, mechanosensory, gustatory, and thermosensory systems^87–92^ (Fig. 7P and Fig. S7D).

Given that Gr5a neurons project to sNPF-sNPF-R-positive circuits near the AL/SOG boundary^37,64^, we propose that taste and pheromonal inputs from Gr5a neurons convey information about male’s sexual experience to the sNPF-sNPF-R circuits, which are processed by the YL neurons. This information is then utilized to generate SELTM via the YL neurons, analogous to the accumulator-memory circuits in the internal clock model for measuring time intervals (Fig. 7Q and Fig. S7E).

## DISCUSSION

In this study, we explore the role of a subset of MB projection neurons, termed ’Yuelao (YL)’ neurons, in the formation of sexual experience-related LTM (SELTM) in male *Drosophila*. These YL neurons express the *orb2* gene, a postsynaptic scaffolding protein crucial for LTM in courtship conditioning (Fig. 1). Our findings demonstrate that sexual experience triggers the formation of LTM for SMD in male flies, which is dependent on the expression of Orb2 in YL neurons (Fig. 2). The molecular mechanisms underlying this LTM involve the RNA-binding protein Fmr1, CREB transcription, and the NMDAR2-CaM-Kinases signaling cascade in YL neurons (Fig. 3 and Fig. 4). Additionally, we uncover that the neuromodulator sNPF in WG neurons and its receptor sNPF-R in YL neurons regulate the release of glutamate, which is essential for LTM formation (Fig. 5). Sexual experience leads to synaptic plasticity and reduces intracellular calcium levels in YL neurons (Fig. 6).

Furthermore, YL neurons exhibit specialization for generating sexual experience-specific appetitive LTM. These neurons receive inputs from taste and pheromonal circuits and project to memory-related brain regions (Fig. 7). Overall, our study provides new insights into the neural circuits and molecular mechanisms underlying taste-related memory formation in *Drosophila*.

Research on *Drosophila* taste memory has predominantly centered on the neural circuitry and molecular mechanisms underlying aversive taste learning. Flies exhibit robust avoidance behaviors after exposure to bitter substances, serving as a warning of potential toxicity. This aversive taste memory is critical for the fly’s survival, enabling it to avoid consuming harmful substances^93^. Although aversive taste LTM has been extensively characterized, the neural circuits mediating appetitive taste LTM remain relatively underexplored^91,94–96^. The SOG is a key brain region involved in processing sweet taste and contact chemoreception^97^, and YL neurons are located above SOG (Fig. 2F). Appetitive taste LTM is crucial for *Drosophila* as it aids in forming associations between food cues and reward, facilitating food selection and foraging behaviors. The discovery of YL neurons is significant as these specialized neurons in the MB might facilitate the formation of taste-related appetitive LTM. Our findings contribute to a comprehensive understanding of the complex relationship between taste and other sensory modalities in memory formation and have broader implications for understanding the neural basis of learning and memory.

Previous studies reported that two pairs of cholinergic MB efferent neurons are required for appetitive LTM retrieval. These MB-V3 cholinergic neurons are specifically recruited for the retrieval of appetitive LTM, but not STM^98,99^. YL neurons are 3 pairs of neurons, and 1 pair of YL neurons are glutaminergic, not cholinergic (Fig. 5O). This single pair of neurons appears solely responsible for SELTM since the knockdown of VGlut, not ChAT, in YL neurons disrupts SELTM (Fig. 5H and Fig. 5L). Thus, we speculate SELTM only requires the function of a single pair of glutaminergic YL neurons, which are functionally distinct from MB-V3 cholinergic neurons. However, the role of these MB-V3 efferent neurons in SELTM has not been investigated in previous studies^98^. This is likely due to the complexity of taste coding and the challenge in isolating taste from other sensory modalities. Our study further investigates the molecular mechanisms by which NMDAR2-CaM-Kinases with sNPF-sNPF-R signaling cascade can jointly modulate Orb2-mediated synaptic protein synthesis.

The sNPF has been identified to play an inhibitory role in certain neural circuits ^100–103^. The inhibitory effects of sNPF are thought to be mediated through the activation of sNPF-R, which leads to the inhibition of neurotransmitter release or the activation of inhibitory neurotransmitter pathways. Specifically, sNPF has been shown to interact with neurotransmitters such as GABA, a primary inhibitory neurotransmitter in the brain, and glutamate, the primary excitatory neurotransmitter ^104^. When sNPF binds to sNPF-R, it can enhance the inhibitory effects of GABA or suppress the excitatory effects of glutamate, leading to a net inhibitory effect on neurotransmission. It has been investigated that Gα_o_ is the crucial regulator of sNPF effects downstream of sNPF-R ^105^ Gα_o_ has been shown to inhibit cAMP production, although it can also have cAMP-independent effects. We showed that sexual experience dramatically decreases calcium in YL neurons (Fig. 6), suggesting that sNPF to sNPF-R circuits in YL neurons act as inhibitory signals. Further molecular analysis with these circuits to understand how this inhibitory modulation occurs will be interesting to expand our understanding of how taste-specific coding leads to LTM for organisms. One of our important findings is that NMDAR2-CaM-Kinases is working together with the sNPF-sNPF-R system to yield taste-related LTM. Since there has been no study about the relationship between sNPF and NMDAR system, this circuit may provide useful toolkits to understand how appetitive taste can induce LTM for organisms.

## CONCLUSION

In conclusion, our study uncovers the pivotal role of the YL neurons in the formation of sexual experience-dependent long-term memory (SELTM) in male *Drosophila*.

Through a series of genetic and pharmacological experiments, we have demonstrated that the activation of YL neurons, which express the Orb2 scaffolding protein, is both necessary and sufficient for the establishment of SELTM. The molecular mechanisms underlying this process involve the intricate interplay of RNA-binding proteins, CREB transcription factors, and the NMDAR2-CaM-Kinases signaling cascade.

Furthermore, our findings reveal the neuromodulatory role of sNPF and its receptor sNPF-R in regulating glutamate release within YL neurons, which is essential for the consolidation of SELTM. The synaptic plasticity and the subsequent reduction in intracellular calcium levels within YL neurons following sexual experience underscore the significance of these neurons in the encoding of appetitive taste memories. Our research not only advances the understanding of the neural and molecular substrates of SELTM but also provides a foundation for future investigations into the complex interplay between experience, memory, and behavior in *Drosophila*. These insights contribute to the broader knowledge of memory formation and could potentially inform studies on learning and memory processes in other organisms, including humans.

## MATERIALS AND METHOD

*Drosophila melanogaster* was cultured under standard laboratory conditions at **25L (for details, see** Fly Stocks and Husbandry). Samples were prepared as described in the METHODS DETAILS. All fly strains are listed in the KEY RESOURCES TABLE.

### Fly Stocks and Husbandry

*Drosophila melanogaster* was raised on cornmeal-yeast medium at similar densities to yield adults with similar body sizes. Flies were kept in 12 h light: 12 h dark cycles (LD) at 25L (ZT 0 is the beginning of the light phase, ZT12 beginning of the dark phase) except for some experimental manipulation (experiments with the flies carrying *tub-GAL80^ts^*). Wild-type flies were Canton-S. To reduce the variation from genetic background, all flies were backcrossed for at least 3 generations to CS strain. All mutants and transgenic lines used here have been described previously.

### Mating Duration Assay

The mating duration assay in this study has been reported ^37,44,106^. To enhance the efficiency of the mating duration assay, we utilized the *Df(1)Exel6234* (DF here after) genetic modified fly line in this study, which harbors a deletion of a specific genomic region that includes the sex peptide receptor (SPR)^107,108^. Previous studies have demonstrated that virgin females of this line exhibit increased receptivity to males^108^. We conducted a comparative analysis between the virgin females of this line and the CS virgin females and found that both groups induced SMD. Consequently, we have elected to employ virgin females from this modified line in all subsequent studies. For naïve males, 40 males from the same strain were placed into a vial with food for 5 days. For single reared males, males of the same strain were collected individually and placed into vials with food for 5 days. For experienced males, 40 males from the same strain were placed into a vial with food for 4 days then 80 DF virgin females were introduced into vials for last 1 day before assay. 40 DF virgin females were collected from bottles and placed into a vial for 5 days. These females provide both sexually experienced partners and mating partners for mating duration assays. At the fifth day after eclosion, males of the appropriate strain and DF virgin females were mildly anaesthetized by CO_2_. After placing a single female into the mating chamber, we inserted a transparent film then placed a single male to the other side of the film in each chamber. After allowing for 1 h of recovery in the mating chamber in 25L incubators, we removed the transparent film and recorded the mating activities. Only those males that succeeded to mate within 1 h were included for analyses. Initiation and completion of copulation were recorded with an accuracy of 10 sec, and total mating duration was calculated for each couple. All assays were performed from noon to 4pm. Genetic controls with *GAL4/+* or *UAS/+* lines were omitted from supplementary figures, as prior data confirm their consistent exhibition of normal LMD and SMD behaviors^37,44,106,109,110^. Hence, genetic controls for LMD and SMD behaviors were incorporated exclusively when assessing novel fly strains that had not previously been examined. In essence, internal controls were predominantly employed in the experiments, as LMD and SMD behaviors exhibit enhanced statistical significance when internally controlled. Within the LMD assay, both group and single conditions function reciprocally as internal controls. A significant distinction between the naïve and single conditions implies that the experimental manipulation does not affect LMD. Conversely, the lack of a significant discrepancy suggests that the manipulation does influence LMD. In the context of SMD experiments, the naïve condition (equivalent to the group condition in the LMD assay) and sexually experienced males act as mutual internal controls for one another. A statistically significant divergence between naïve and experienced males indicates that the experimental procedure does not alter SMD. Conversely, the absence of a statistically significant difference suggests that the manipulation does impact SMD. Hence, we incorporated supplementary genetic control experiments solely if they deemed indispensable for testing. All assays were performed from noon to 4 PM. We conducted blinded studies for every test^111,112^.

### Pharmacology

*Drosophila* were subjected to treatment with the NMDA receptor antagonist MK-801, also known as Dizocilpine. The MD assays were conducted following acute administration of MK-801 to the flies (over a period of 24 hours) at a concentration of 0.3 mM, incorporated into their dietary regimen. The selected drug concentration was informed by findings from a preceding study ^113^. For the control cohorts, an equal volume of the vehicle solution, which in this case was distilled water, was mixed into the fly food.

### T-maze Aversive Conditioning Protocol

The T-maze was a custom-made 3D printed arena (polyoxymethylene material) adapted and simplified from the conventional Tully T-maze ^114^. The arena was modified to ensure least interference from the reflected light. Innate preference was determined by testing the choice of a group of naive flies (50-60 in number, 3–5 days old) to the odors and colors against paraffin oil and darkness, respectively. For the training protocol, a different group of 50-60 flies was aspirated into the training tube that was lined with an embedded copper grid. Aversive reinforcement of the stimuli (CS+: odor/light/bimodal) was provided for 1 min paired with AC 90V electric shock. One training trial consisted of 15 shock pulses, each pulse lasting 1.8 s followed by an interval of 3 s. After training, flies were given 1 min of clean air or kept in darkness, then followed by 1 min of testing (odor OTC and MCH vs paraffin oil/light vs dark). There was no electric shock presented during the testing. For bimodal training, CS + constituted a simultaneous presentation of the light, odor and shock, were also performed to rule out non-associative effects. All experiments were conducted at the same time-window for all days to control for circadian variation in wavelength preferences. Suction was always provided to prevent odor contamination or accumulation in the center of the maze. The tube lined with the copper grid was used in both the training and the testing phase to reproduce the same contexts. All training and testing were done at 25°C and 70% relative humidity in a dark arena. Flies inside the arena were handled only in red light (>720 nm). Different groups of flies were used to observe innate and learned behavior. Preference indices were calculated for both using the following formula:

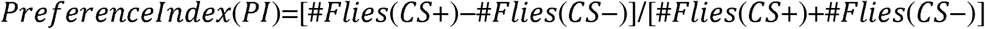

In experiments where long-lasting memory retrieval was necessary, trained flies were collected and stored in vials with food at 25°C and darkness for specified time.

### Immunostaining

The dissection and immunostaining protocols for the experiments are described elsewhere ^106^. After 5 days of eclosion, the *Drosophila* brain was taken from adult flies and fixed in 4% formaldehyde at room temperature for 30 minutes. The sample was washed three times (5 minutes each) in 1% PBT and then blocked in 5% normal goat serum for 30 minutes. The sample next be incubated overnight at 4℃ with primary antibodies in 1% PBT, followed by the addition of fluorophore-conjugated secondary antibodies for one hour at room temperature. The brain was mounted on plates with an antifade mounting solution (Invitrogen) for imaging purposes.

Samples were imaged with Zeiss LSM880. Antibodies were used at the following dilutions: Chicken anti-GFP (1:500, Invitrogen), mouse anti-nc82 (1:50, DSHB), rabbit anti-DsRed (1:500, Rockland Immunochemicals), Alexa-488 donkey anti-chicken (1:200, Jackson ImmunoResearch), Alexa-555 goat anti-rabbit (1:200, Invitrogen), Alexa-647 goat anti-mouse (1:200, Jackson ImmunoResearch). For specific antibodies used for immunostaining, see **KEY RESOURCE TABLES**.

### Quantitative Analysis of Fluorescence Intensity

To ascertain calcium levels and synaptic intensity from microscopic images, we dissected and imaged five-day-old flies of various social conditions and genotypes under uniform conditions. The GFP signal in the brains and VNCs was amplified through immunostaining with chicken anti-GFP primary antibody. Image analysis was conducted using ImageJ software. For the quantification of fluorescence intensities, an investigator, blinded to the fly’s genotype, thresholded the sum of all pixel intensities within a sub-stack to optimize the signal-to-noise ratio, following established methods ^115^. The total fluorescent area or region of interest (ROI) was then quantified using ImageJ, as previously reported. For CaLexA signal quantification, we adhered to protocols detailed by Kayser et al.^116^, which involve measuring the ROI’s GFP-labeled area by summing pixel values across the image stack. This method assumes that changes in the GFP-labeled area are indicative of alterations in the CaLexA signal, reflecting synaptic activity. ROI intensities were background-corrected by measuring and subtracting the fluorescent intensity from a non-specific adjacent area, as per Kayser et al. ^116^. For the analysis of GRASP or tGRASP signals, a sub-stack encompassing all synaptic puncta was thresholded by a genotype-blinded investigator to achieve the optimal signal-to-noise ratio. The fluorescence area or ROI for each region was quantified using ImageJ, employing a similar approach to that used for CaLexA quantification^115^.

### Colocalization Analysis

Before the colocalization analysis, an investigator, blinded to the fly’s genotype, thresholded the sum of all pixel intensities within a sub-stack to optimize the signal-to-noise ratio, following established methods ^115^. To perform colocalization analysis of multi-color fluorescence microscopy images in this study, we employed ImageJ software ^117^. In brief, we merged image channels to form a composite with accurate color representation and applied a threshold to isolate yellow pixels, signifying colocalization. The measured “area” values represented the colocalization zones between fluorophores. To determine the colocalization percentage relative to the total area of interest (e.g., GFP or RFP), we adjusted thresholds to capture the full fluorophore areas and remeasured to obtain total areas. The colocalization efficiency was calculated by dividing the colocalized area by the total fluorophore area. All samples were imaged uniformly.

### Single-nucleus RNA-sequencing Analyses

The snRNAseq dataset analyzed in this paper has been published^45^ and available at the Nextflow pipelines (VSN, https://github.com/vib-singlecell-nf), the availability of raw and processed datasets for users to explore, and the development of a crowd-annotation platform with voting, comments, and references through SCope (https://flycellatlas.org/scope), linked to an online analysis platform in ASAP (https://asap.epfl.ch/fca). For the generation of the tSNE plots, we utilized the Fly SCope website (https://scope.aertslab.org/#/FlyCellAtlas/*/welcome). Within the session interface, we selected the appropriate tissues and configured the parameters as follows: ’Log transform’ enabled, ’CPM normalize’ enabled, ’Expression-based plotting’ enabled, ’Show labels’ enabled, ’Dissociate viewers’ enabled, and both ’Point size’ and ’Point alpha level’ set to maximum. For all tissues, we referred to the individual tissue sessions within the ’10X Cross-tissue’ RNAseq dataset. Each tSNE visualization depicts the coexpression patterns of genes, with each color corresponding to the genes listed on the left, right, and bottom of the plot. The tissue name, as referenced on the Fly SCope website is indicated in the upper left corner of the tSNE plot. Dashed lines denote the significant overlap of cell populations annotated by the respective genes. Coexpression between genes or annotated tissues is visually represented by differentially colored cell populations. For instance, yellow cells indicate the coexpression of a gene (or annotated tissue) with red color and another gene (or annotated tissue) with green color. Cyan cells signify coexpression between green and blue, purple cells for red and blue, and white cells for the coexpression of all three colors (red, green, and blue). Consistency in the tSNE plot visualization is preserved across all figures.

Single-cell RNA sequencing (scRNA-seq) data from the *Drosophila melanogaster* were obtained from the Fly Cell Atlas website (https://doi.org/10.1126/science.abk2432). Oenocytes gene expression analysis employed UMI (Unique Molecular Identifier) data extracted from the 10x VSN oenocyte (Stringent) loom and h5ad file, encompassing a total of 506,660 cells. The Seurat (v4.2.2) package (https://doi.org/10.1016/j.cell.2021.04.048) was utilized for data analysis. Violin plots were generated using the “Vlnplot” function, the cell types are split by FCA.

### Statistical Tests

Statistical analysis of mating duration assay was described previously^37,44,106^. More than 50 males (naïve, experienced and single) were used for mating duration assay. Our experience suggests that the relative mating duration differences between naïve and experienced condition and singly reared are always consistent; however, both absolute values and the magnitude of the difference in each strain can vary. So, we always include internal controls for each treatment as suggested by previous studies ^118^. Therefore, statistical comparisons were made between groups that were naïvely reared, sexually experienced and singly reared within each experiment. As mating duration of males showed normal distribution (Kolmogorov-Smirnov tests, p > 0.05), we used two-sided Student’s t tests. The mean ± standard error (s.e.m) (*\*\*\*\* = p < 0.0001, *** = p < 0.001, ** = p < 0.01, * = p < 0.05*). All analysis was done in GraphPad (Prism). Individual tests and significance are detailed in figure legends. Besides traditional *t*-test for statistical analysis, we added estimation statistics for all MD assays and two group comparing graphs. In short, ‘estimation statistics’ is a simple framework that—while avoiding the pitfalls of significance testing—uses familiar statistical concepts: means, mean differences, and error bars. More importantly, it focuses on the effect size of one’s experiment/intervention, as opposed to significance testing ^119^. In comparison to typical NHST plots, estimation graphics have the following five significant advantages such as (1) avoid false dichotomy, (2) display all observed values (3) visualize estimate precision (4) show mean difference distribution. And most importantly (5) by focusing attention on an effect size, the difference diagram encourages quantitative reasoning about the system under study ^120^. Thus, we conducted a reanalysis of all our two group data sets using both standard *t* tests and estimate statistics. In 2019, the Society for Neuroscience journal eNeuro instituted a policy recommending the use of estimation graphics as the preferred method for data presentation ^121^.

### Courtship Conditioning/Courtship Memory Test and Analysis

Animals were prepared for courtship conditioning as previously described^122^. In short, males of appropriate strain were collected during the pupal stage and individually housed in clean glass tubes containing fresh food for 4 days post-eclosion to prevent pre-test social interaction. To prevent anesthetic effects, we used a mouth aspirator to transfer or collect animals. Trainer females were prepared by housing day 3–5 virgin Canton-S females with Canton-S males in food-containing glass tubes overnight.

Immobilized tester females were prepared by decapitating day 4–5 virgin Canton-S females with fine scissors under CO_2_ anesthesia. Protocols for long-term courtship conditioning followed those described previously ^123^, with some modifications.

Briefly, a 4-day old male was paired with a mated Canton-S female for 5 h in the food chamber (2.0 ml tube). Trained (or sham-trained) males were individually housed in a fresh food tube for 24 h prior to testing, whereupon courtship activity of a trained (or sham-trained) male toward an immobilized (decapitated) tester female was recorded for 10 min using a high-frame digital camcorder (60hz). All assays were manually scored for courtship index, blind to the genotype and to the extent possible, the experimental condition. Positive contributions to courtship index (CI), defined as the percentage of time a male performs courtship behavior over a 10 min interval, included all courtship activities such as orienting, tapping, singing, licking, and attempting to copulate. Courtship memory was quantified through computation of a memory performance index (MPI), which is the relative reduction of CI of individual trained male (^i^CI_T_) from the mean CI of sham-trained males (^m^CI_S_): MPI= (^m^CI_S_-^i^CI_T_)/ ^m^CI_S._

### Generation of Transgenic Flies

To create a membrane-anchored version of *UAS-t-sNPF* strains, we employed the previously described GPI-linker technique to produce a sNPF variant that is attached to the cell membrane ^124^. Using the aforementioned approaches, we created several transgenic flies with *UAS-t-sNPF*. The peptide sequence expressed by the above gene is:

*UAS-t-sNPF-1-ML*: MSALLILALVGAAVAAQRSPSLRLRFGNEQKLISEEDLGNGAGFATPVTLALVP

ALLATFWSLL

*UAS-t-sNPF-1-SEC*: MSALLILALVGAAVAAQRSPSLRLRFGNEQKLISEEDLGN *UAS-t-sNPF-2-ML:* MSALLILALVGAAVAWFGDVNQKPIRSPSLRLRF GNEQKLISEEDLGNGAGFATPVTLALVPALLATFWSLL

*UAS-t-sNPF-2-SEC:*

MSALLILALVGAAVAWFGDVNQKPIRSPSLRLRFGNEQKLISEEDLGN

*UAS-t-sNPF-2_12-19_-SEC:* MSALLILALVGAAVASPSLRLRFGNEQKLISEEDLGN *UAS-t-sNPF-3-ML*: MSALLILALVGAAVAKPQRLRWGNEQKLISEEDLGNGAG FATPVTLALVPALLATFWSLL

*UAS-t-sNPF-3-SEC:* MSALLILALVGAAVAKPQRLRWGNEQKLISEEDLGN

Synthesis of DNA, subcloning into the pUAST-attB vector, and the introduction of transgenes into the *Drosophila* genome via microinjection services were performed by Qidong Fungene Biotechnology Co., Ltd. (http://www.fungene.tech).

## KEY RESOURCES TABLE

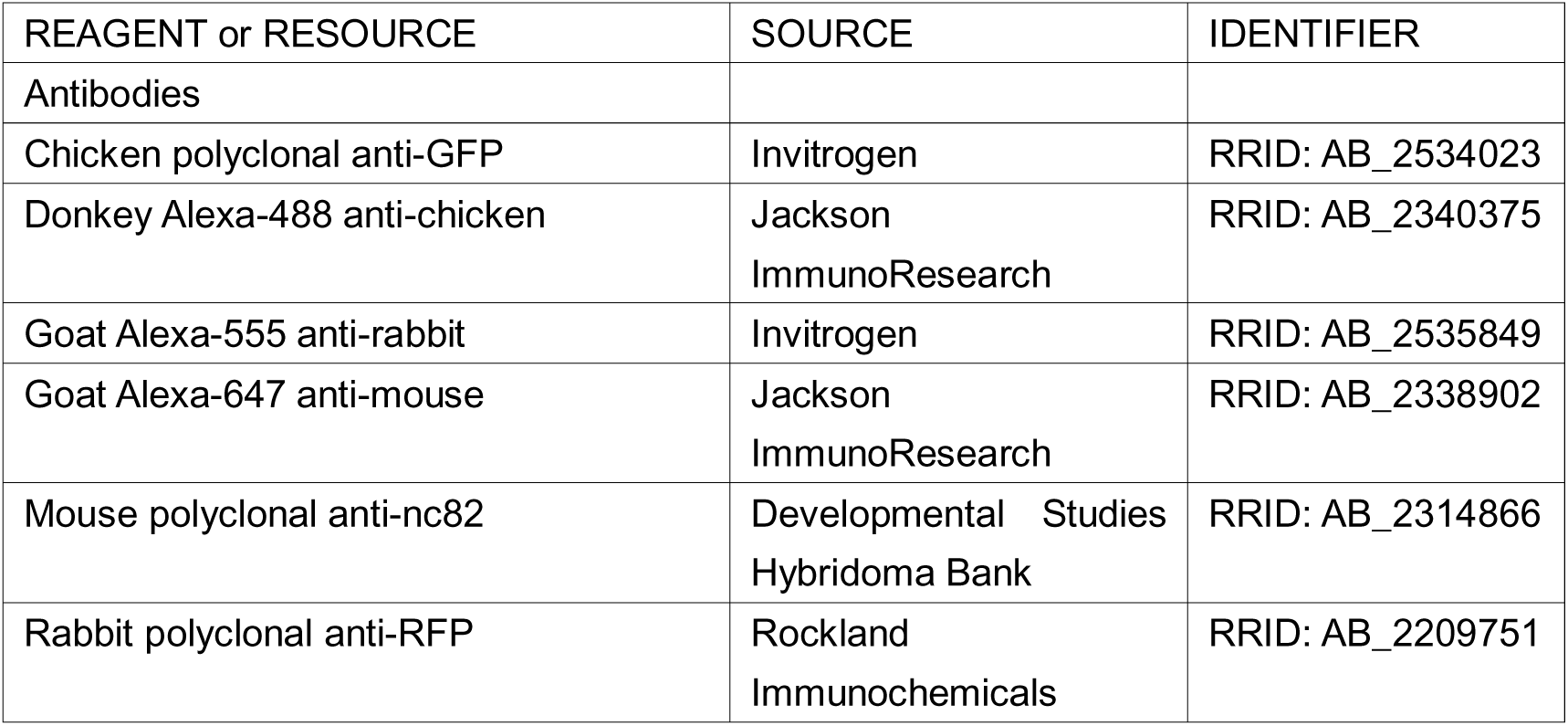
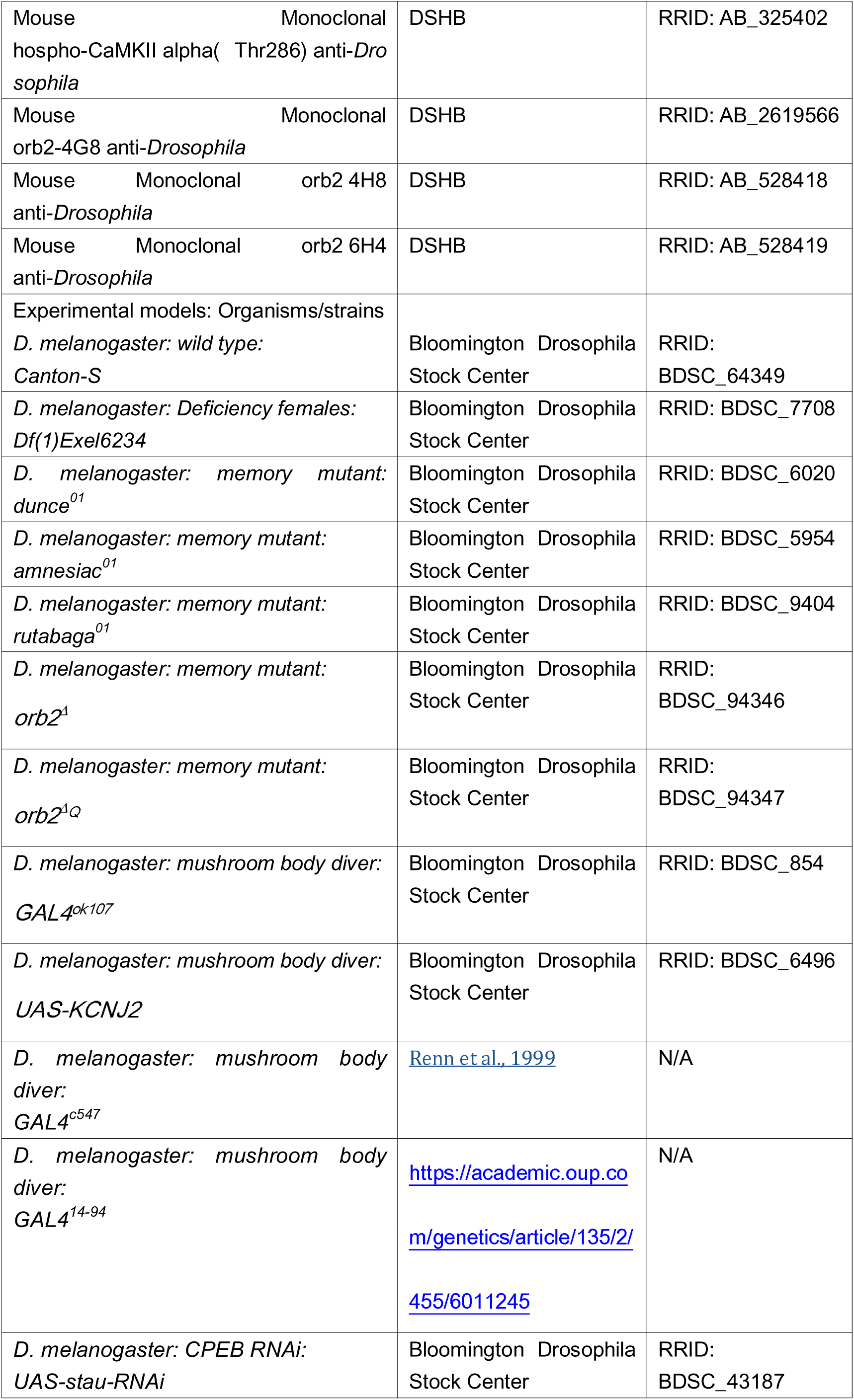
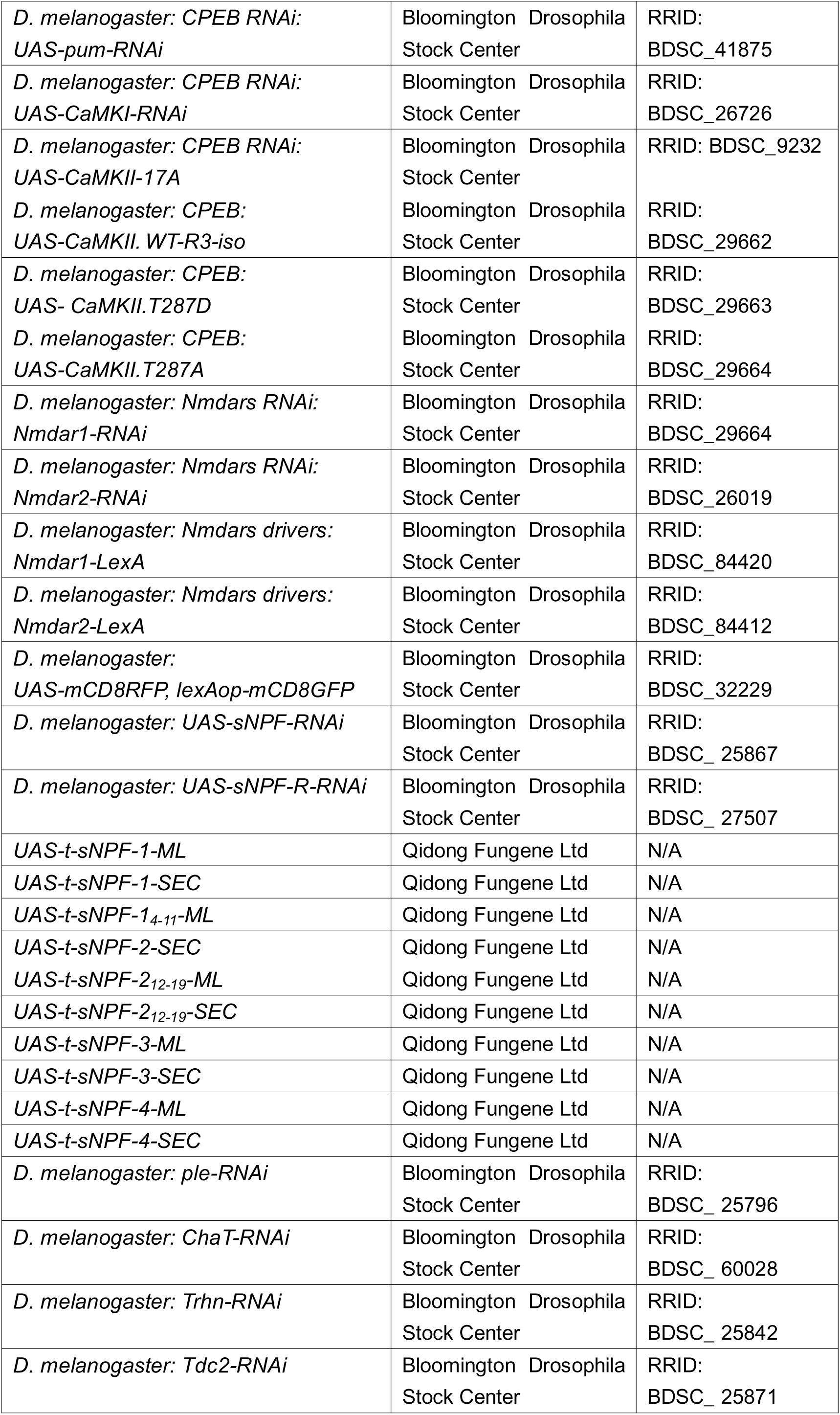
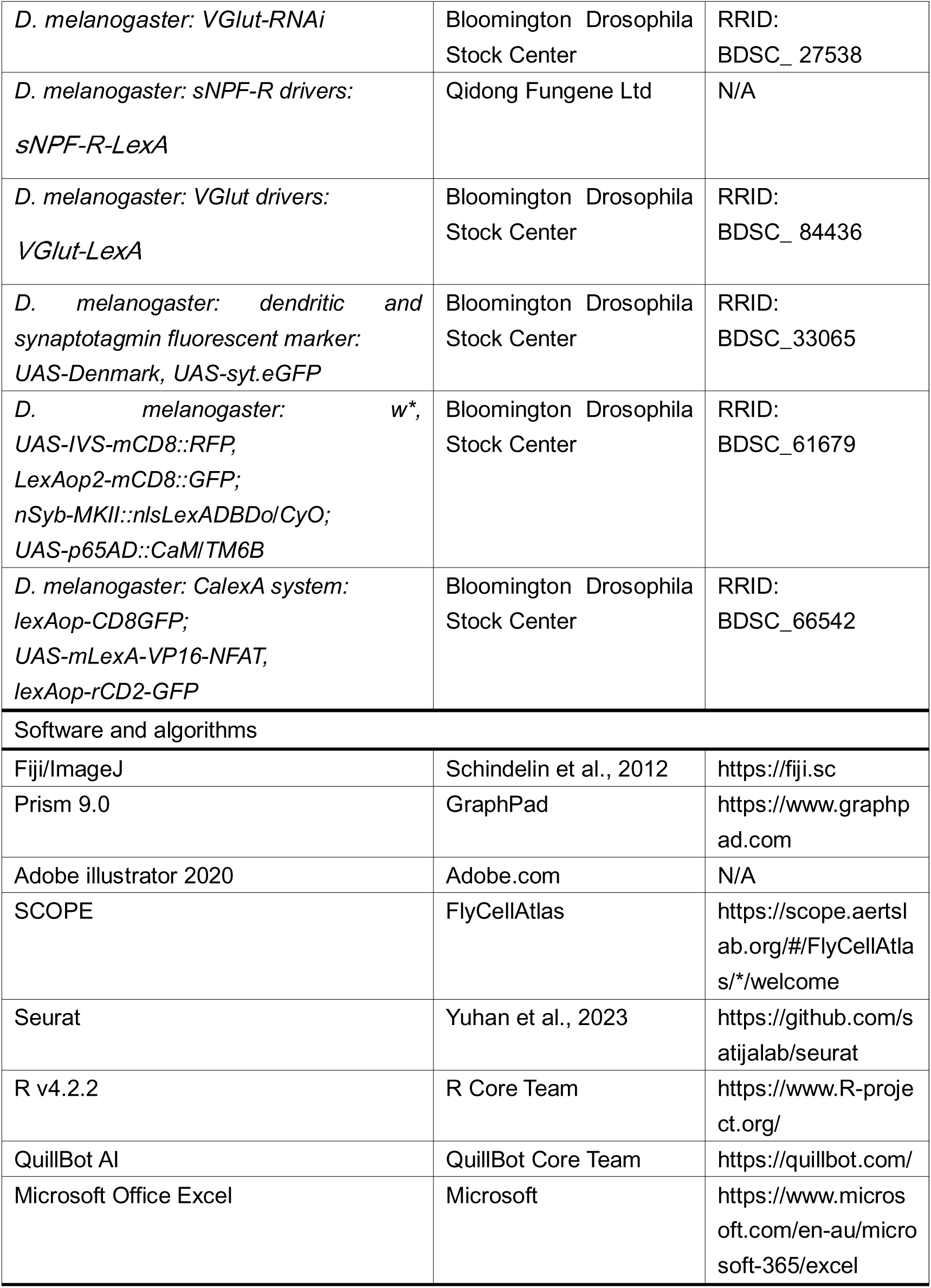

## RESOURCE AVAILABILITY

### Lead contact

Further information and requests for resources and reagents should be directed to and will be fulfilled by the lead contact, Woo Jae Kim.

### Data and code availability

- All data reported in this paper will be shared by the lead contact upon request.
- This paper does not report original code. The URL of the codes used in this paper are listed in the key resources table.
- Any additional information required to reanalyze the data in this paper is available from the lead contact upon request.

## Acknowledgments

We thank Drs. Yuh Nung Jan and Lily Yeh Jan (UCSF, USA) for helpful comments and support on this paper. We are also very appreciative to the colleagues who supplied us with several fly strains: Drs. Young-Joon Kim and Sung-Eun Yoon (Korea Drosophila Resource Center, KDRC). This research was supported a University of Ottawa Startup grant 602496 to WJK, Startup funds from HIT Center for Life Science to WJK, a University of Ottawa Interdisciplinary Research Group Funding Opportunity (IRGFO stream 1 and 2) grants 148101 and 148747 to WJK, a Natural Sciences and Engineering Research Council of Canada (NSERC) Discovery grant (reference: 211406) to WJK, a University of Ottawa Brain and Mind Research Institute/Center for Neural Dynamics Open call project grant 150950 to WJK, a Mitacs Globalink Research Internship Program grant 17268 to WJK. This research was also supported by the Brain Pool Program of the National Research Foundation in Korea grant ZYM5041911 to WJK, Burroughs Wellcome Fund Collaborative Research Travel Grants (reference: 1017486) to WJK and a NVIDIA Academic Hardware Grant Program to WJK. The funders had no role in study design, data collection and analysis, decision to publish, or preparation of the manuscript.

## DECLARATION OF GENERATIVE AI AND AI-ASSISTED TECHNOLOGIES IN THE WRITING PROCESS

During the creation of this work, the author(s) utilized Moonshot AI https://kimi.moonshot.cn/ to rephrase English sentences, verify English grammar, and detect plagiarism, as none of the authors of this paper are native English speakers.

After using this tool/service, the author(s) reviewed and edited the content as needed and take(s) full responsibility for the content of the publication.

## Author contributions

Conceptualization: Woo Jae Kim.

Data curation: Dongyu Sun, Xiaoli Zhang, Hongyu Miao, Tianmu Zhang, and Woo Jae Kim.

Formal analysis: Dongyu Sun, Xiaoli Zhang, Hongyu Miao, Tianmu Zhang, and Woo Jae Kim.

Funding acquisition: Woo Jae Kim.

Investigation: Woo Jae Kim.

Methodology: Woo Jae Kim.

Project administration: Woo Jae Kim.

Resources: Woo Jae Kim.

Supervision: Woo Jae Kim.

Validation: Dongyu Sun, Xiaoli Zhang, Hongyu Miao, Tianmu Zhang, and Woo Jae Kim.

Visualization: Woo Jae Kim.

Writing – original draft: Woo Jae Kim.

Writing – review & editing: Dongyu Sun and Woo Jae Kim.

## Supplementary information

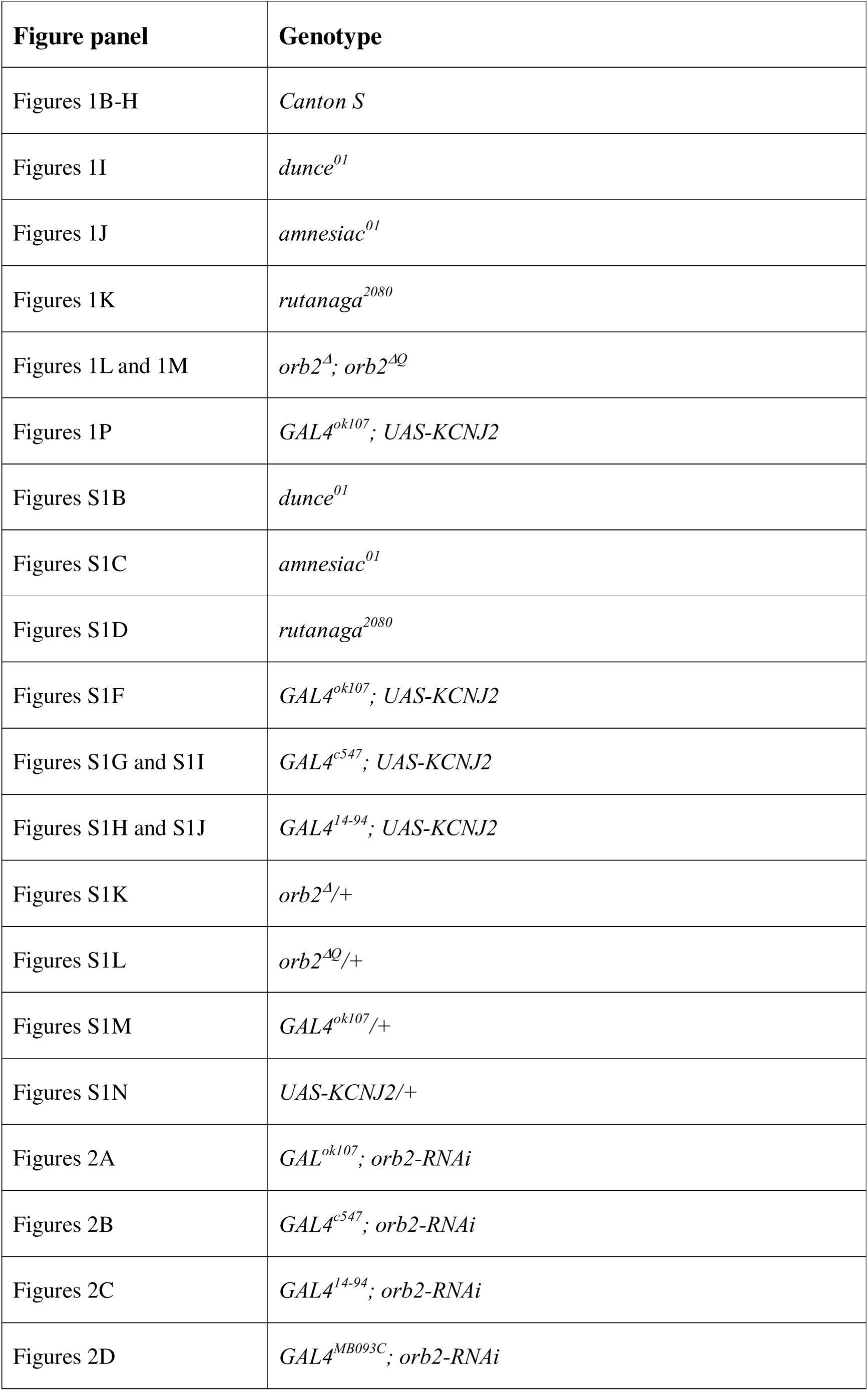
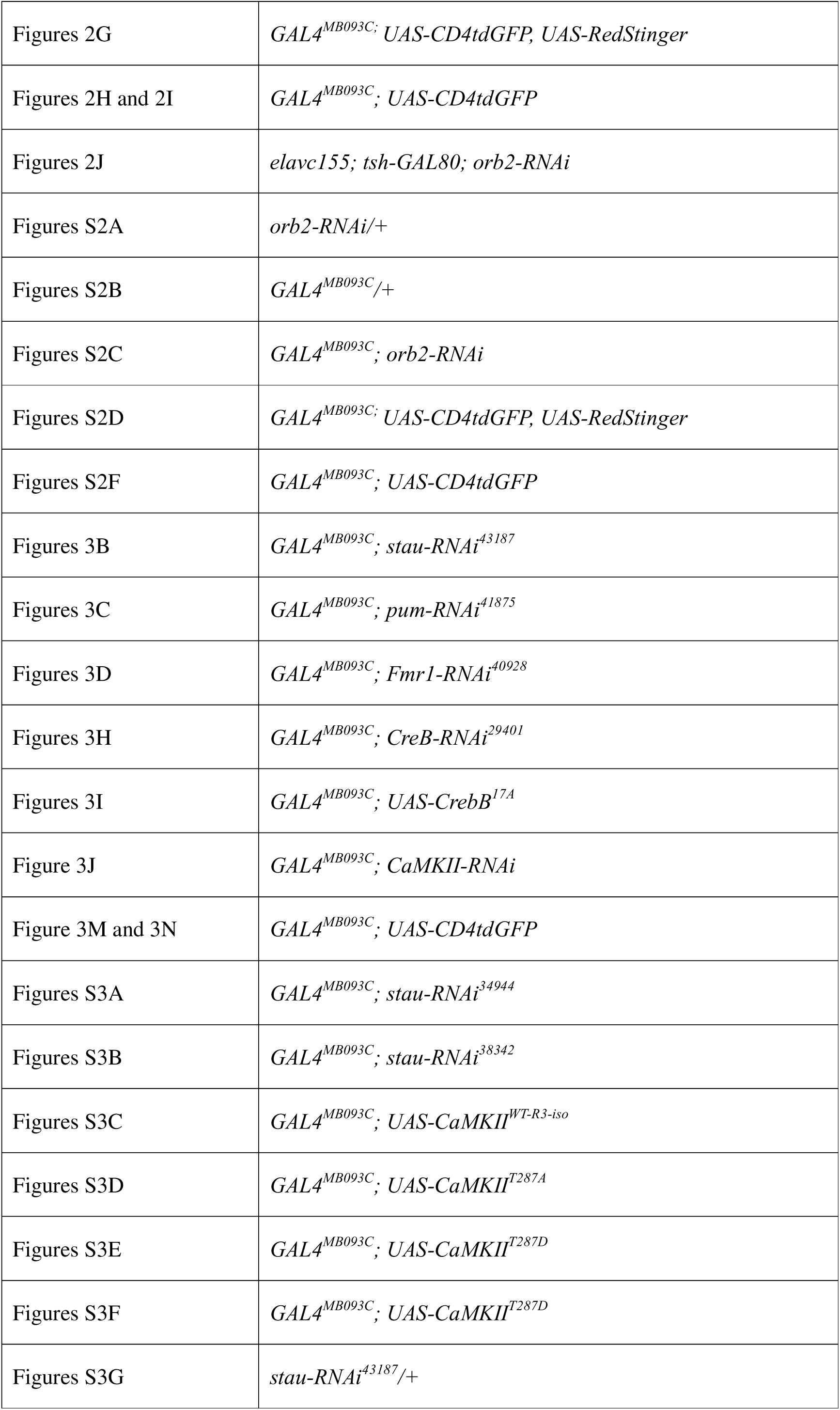
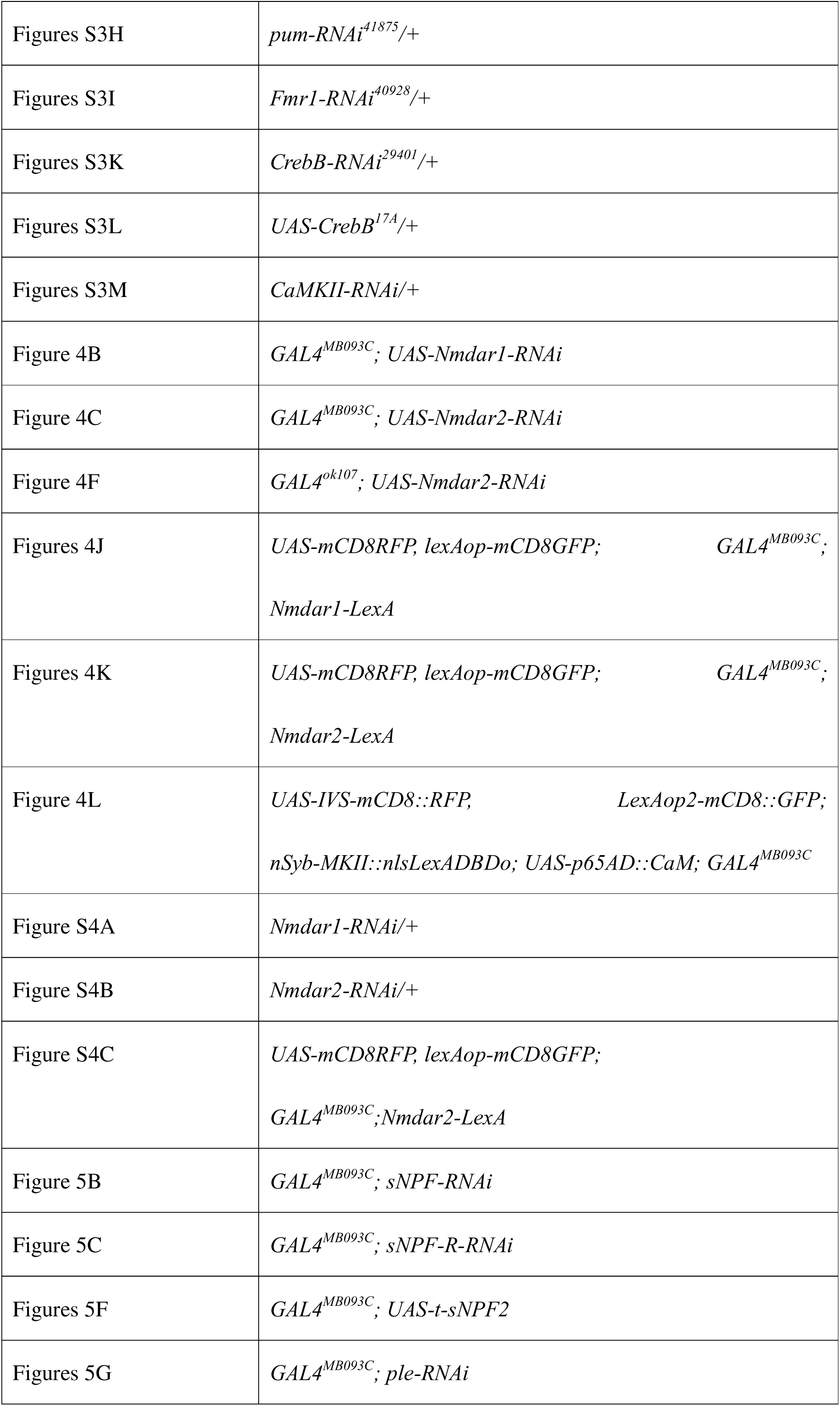
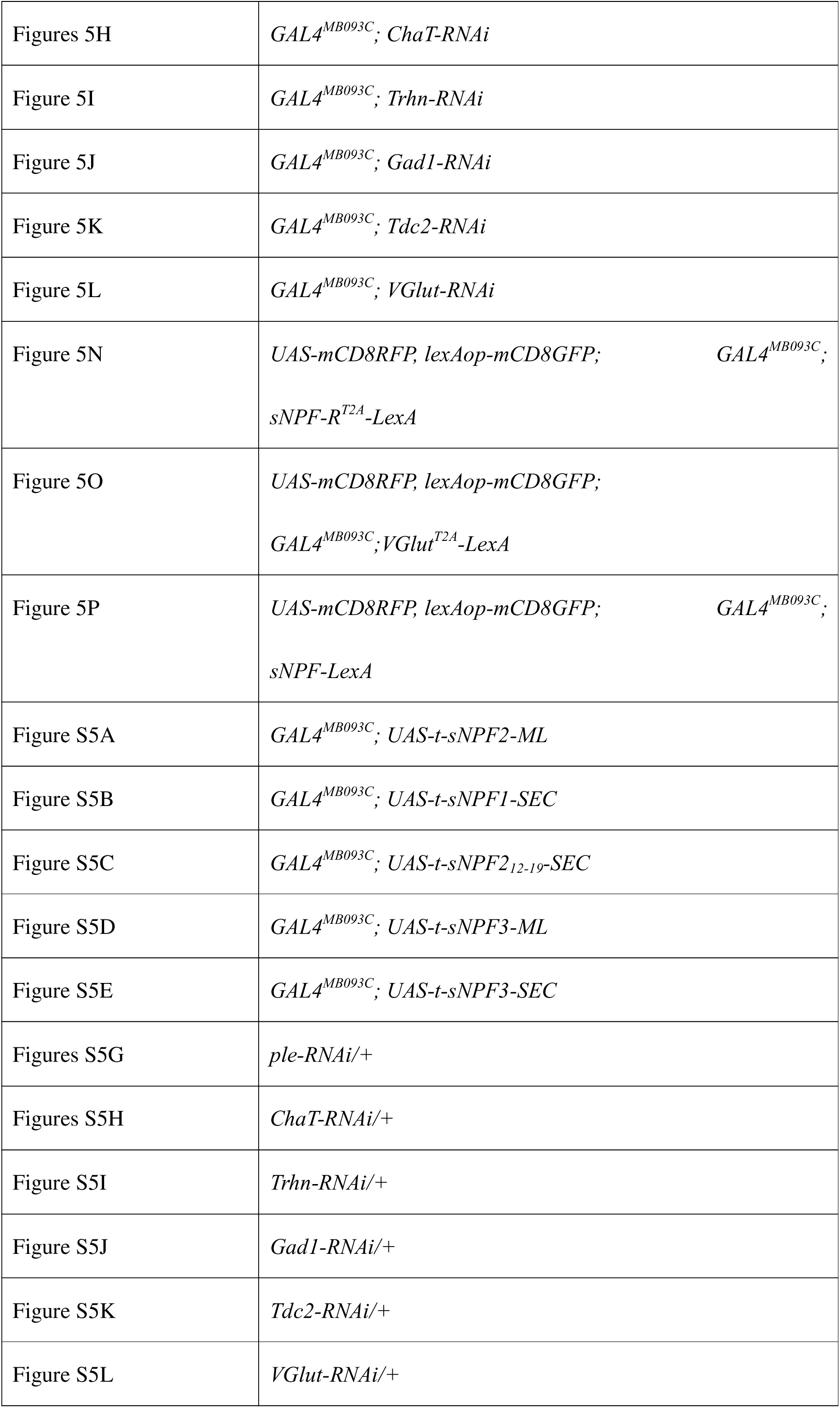
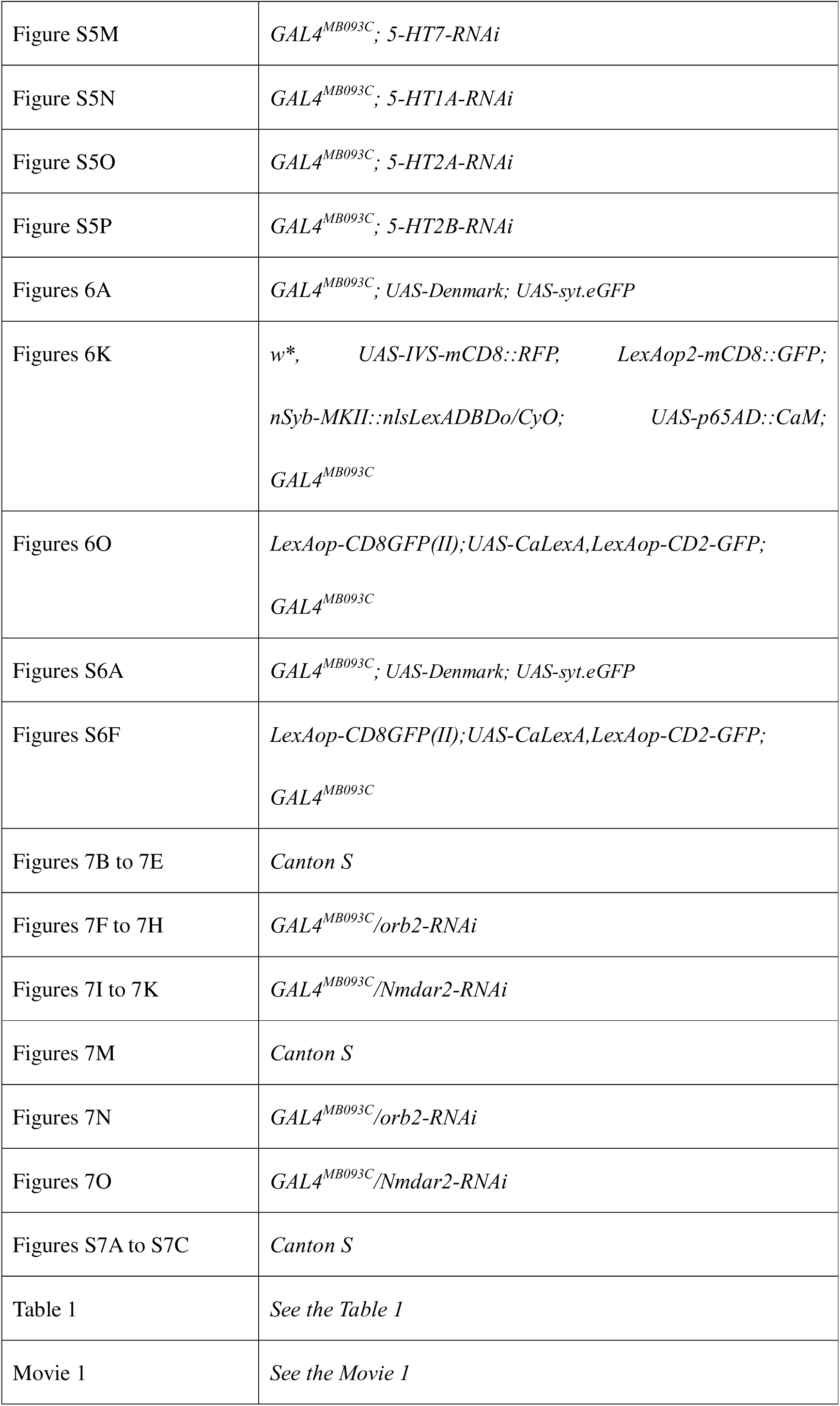

Fig S1. The extended mating duration behavior elicited by rival (LMD behavior) does not depend on the function of Orb2 within the MB.

(A) The schematic diagram of interval timing simulated based on relevant literature sources shows that when sexual experience is transmitted as information, CLK cells are “pacemakers,” Gr5a cells are “mode switches,” and sNPF cells are “accumulators.” The relevant information is integrated into “memory,” and YL neurons are the key to the “memory” element. Finally, the information is handed over to the downstream “decision” part to realize the relevant behavior regulation.

(B-D) LMD assays for Memory-defective mutant fruit flies, *dunce^01^*(B)*, amnesiac^01^* (C) *and rutabaga^2080^* (D).

(E) Brain and VNC of male flies expressing *orb2* with *GFP,* were immunostained with anti-GFP (green) and nc82 (magenta) antibodies. Scale bars represent 100 μm. Data from Kruttner, S. et al. ^33^.

(F-J) LMD assays for *GAL4^ok107^* (F) *GAL4**^c547^*** (G) and *GAL4**^14-94^*** (H) mediated electrical silencing *via UAS-KCNJ2*.and SMD assays for *GAL4**^c547^*** (I) and *GAL4**^14-94^***

(J) mediated electrical silencing *via UAS-KCNJ2*.

(K-N) SMD assays for wide type background with *orb2^D^/+* (K) *orb2^DQ^/+* (L) *GAL4^ok107^/+* (M) and *UAS-KCNJ2/+* (N).

Fig S2. Orb2 is necessary for the manifestation of SMD behavior in YL neurons and Orb2 expression.

(A-C) SMD assays for wide type background with *orb2-RNAi* (A) *and GAL4^MB093C^* (B) and SMD assays for *GAL4^MB093C^* mediated knockdown of Orb2 via *orb2-RNAi* (C).

(D) Brain and VNC of male flies expressing *GAL4^MB093C^* with *UAS-CsChrimson-mVenus,* were immunostained with anti-GFP (green) and nc82 (magenta) antibodies. Scale bars represent 100 μm. Boxes indicate the magnified regions to clearly show the expression patterns of neurons in brain labeled by *GAL4^MB093C^* driver. The data from Flylight (https://www.janelia.org/project-team/).

(E) Quantification was performed for the GFP fluorescence in brain and VNC between naïve and experienced male flies. The GFP fluorescence was normalized relative to the fluorescence of the relatively with GFP. The conditions of flies are described above: naïve, naïve male flies; exp., male flies with sexual experience. Bars represent the mean of the normalized GFP fluorescence level with error bars representing the SEM. Asterisks represent significant differences, as revealed by the Student’s *t* test and ns represents non-significant difference (*\*p < 0.05, **p < 0.01, ***p < 0.001*). See the METHODS for a detailed description of the description of the colocalization analysis used in this study.

(F) MB-α/β-lobe of male flies expressing *GAL4^MB093C^* with *UAS-CD4tdGFP* were immunostained with anti-GFP (green) and anti-orb2^6H4^ (red). Just show the MB-α/β-lobe of male flies is clearly show the expression patterns of neurons in brain labeled by *GAL4^MB093C^* driver and see the Significant differences in MB-α/β-lobe.

(G-J) Colocalization analysis of GFP and DsRed staining, normalized to total GFP and DsRed areas. Bars represent the mean GFP (green column) and DsRed (red column) See the METHODS for a detailed description of the colocalization analysis used in this study.

Fig S3. Not all of the CPEB in YL neuron is necessary for the manifestation of SMD behavior.

(A-B) SMD assays for *GAL4^MB093C^* mediated knockdown of stau and pum via *stau-RNAi^34944^*(A) and *pum -RNAi^38342^*(B).

(C) SMD assays for *GAL4^MB093C^* mediated knockdown of CaMKI via *CaMKI-RNAi*.

(D) SMD assays for *GAL4^MB093C^* overexpress CaMKII via *UAS-CamKII^WT-R3-iso^* .

(E-F) SMD assays for *GAL4^MB093C^* express CaMKII mutant via *UAS-CaMKII^T287A^* (E) *and UAS-CaMKII^T287D^*(F).

(G-L) SMD assays for wide type background with *stau-RNAi* (G), *pum-RNAi* (H)*, Fmr1-RNAi* (I), CrebB*-RNAi^29401^* (J), *UAS-CrebB^17A^* (K) and *CaMKII-RNAi* (L).

Fig S4. Localization of NMDAR2 in YL neurons and neuronal response of YL neurons following reception of mating cues.

(A-B) SMD assays for wide type background with *Nmdar1-RNAi* (A), and *Nmdar2-RNAi* (B).

(C) Brain of female flies expressing *Nmdar2-LexA* and *GAL4^MB093C^* together with *CD4tdGFP-LexAop, UAS-Redstinger* were immunostained with anti-GFP (green), anti-DsRed (red), and nc82 (blue) antibodies. Scale bars represent 100 μm. Boxes indicate the magnified regions of interest presented in the left two panels are presented as a grey scale to clearly show the expression patterns of neurons in brain labeled by *GAL4^MB093C^* driver and the right two panels are zoom in the boxes indicate the magnified regions to clearly shows the overlapping.

(C-F) GFP is pseudo-colored as “red hot”. Indicate the magnified regions of interest presented in the left panels. The left three panels are presented as a grey scale to clearly show the expression patterns of neurons in brain labeled by *GAL4^MB093C^* driver.

And zoom in the indicate the magnified regions, MB-α/β-lobe (D-E). SOG(G). And quantification was performed for the GFP fluorescence in MB-α/β-lobe (F) and SOG

(H) between naïve and experienced male flies. The GFP fluorescence was normalized relative to the fluorescence of the nc82. The conditions of flies are described above: naïve, naïve male flies; exp., male flies with sexual experience. Bars represent the mean of the normalized GFP fluorescence level with error bars representing the SEM. Asterisks represent significant differences, as revealed by the Student’s *t* test and ns represents non-significant difference. For detailed methods, see the METHODS for a detailed description of the immunostaining procedure used in this study.

Fig S5. YL neurons are sNPFR cells but not serotonergic neurons

(A-E) SMD assays for *GAL4^MB093C^* expression for self-coupling of sNPF and *UAS-t-sNPF2-ML* (A), *UAS-sNPF1-SEC* (B), *UAS-sNPF2_12-19_-SEC* (C), *UAS-sNPF3-ML* (D) and *UAS-sNPF3-SEC* (E).

(F-L) SMD assays for wide type background with *UAS-t-sNPF2* (F), *ple-RNAi* (G), *ChaT-RNAi* (H), *Trhn-RNAi* (I), *Gad1-RNAi* (J), *Tdc2-RNAi* (K) and *VGlut-RNAi* (L).

(M-P) SMD assays for *GAL4^MB093C^* mediated knockdown of 5HT7, 5HT1A, 5HT2A and 5HT2B via *5HT7-RNAi* (M), *5HT1A-RNAi* (N), *5HT2A-RNAi* (O) and *5HT2B-RNAi* (P).

(Q) Illustration depicting the local peptidergic transmission of WG neurons and YL neurons, The red dots symbolize the binding of glutamate to red *Nmdar2*, while the black dots symbolize the binding of sNPF to black sNPF-R, which controls YL neurons and releases glutamate to regulate neurons further down the line.

Fig S6. Alterations in the calcium levels inside YL neurons following sexual encounters.

(A-C) Quantification of brian, which is formed by *GAL4^MB093C^* drivers together with *UAS-Denmark* and *UAS-syteGFP* (also see at Figure 6A) in brain (A), between naïve and experienced male flies. The possible synaptic interactions in naïve and experienced male flies. The six small panels are presented as a red scale to show the GFP and DsRed fluorescence marked by threshold function of ImageJ. And quantification was performed for the GFP and DsRed fluorescence in these regions between naïve and experienced male flies. The GFP and DsRed fluorescence was normalized relative to the fluorescence of the nc82. The conditions of flies are described above: naïve, naïve male flies; exp., male flies with sexual experience. Bars represent the mean of the normalized GFP and DsRed fluorescence level with error bars representing the SEM. Asterisks represent significant differences, as revealed by the Student’s *t* test and ns represents non-significant difference (*\*p < 0.05, **p < 0.01, ***p < 0.001*). See the METHODS for a detailed description of the fluorescence intensity analysis used in this study.

(D-E) Quantification of brain, which is formed by *GAL4^MB093C^*, between naïve and experienced male flies (also see Figure 6 K-N). The possible synaptic interactions in naïve and experienced male flies. The small panels are presented as a red scale to show the GFP fluorescence marked by threshold function of ImageJ. And quantification was performed for the GFP fluorescence in these regions between naïve and experienced male flies. The GFP fluorescence was normalized relative to the fluorescence of the nc82. The conditions of flies are described above: naïve, naïve male flies; exp., male flies with sexual experience. Bars represent the mean of the normalized GFP fluorescence level.

(F) Different levels of neural activity of the brian as revealed by the CaLexA system in naïve, single and experienced flies. Male flies expressing *GAL4^MB093C^* along with *LexAop-CD2-GFP, UAS-mLexA-VP16-NFAT and LexAop-CD8-GFP-A2-CD8-GFP* were dissected after 5 days of growth (mated male flies had 1-day of sexual experience with virgin females). The dissected brains were then immunostained with anti-GFP (fire) and anti-nc82 (grey). GFP is pseudo-colored as “red hot”. Boxes indicate the magnified regions of interest presented in the bottom panels. Scale bars represent 100 μm.

(G-H) Quantification was performed for the GFP fluorescence in brain between naïve and experienced male flies. The GFP fluorescence was normalized relative to the fluorescence of the nc82.The conditions of flies are described above: naïve, naïve male flies; exp., male flies with sexual experience. Bars represent the mean of the normalized GFP fluorescence level with error bars representing the SEM. Asterisks represent significant differences, as revealed by the Student’s *t* test and ns represents non-significant difference (*\*p < 0.05, **p < 0.01, ***p < 0.001*). See the +METHODS for a detailed description of the fluorescence intensity analysis used in this study.

(Q) Upon receiving the sexual experience signal, the sNPF-R and NMDAR2 receptors are activated by Glu and sNPF, respectively. This triggers a sequence of events in YL cells, where CPEB proteins oligomerize and CaMKI and CaMKII phosphorylation regulation to build the memory of SMD behavior. The specific CPEB protein is depicted in the image. Subsequently, Glu transforms the memories into data and transmits it to the prospective downstream neurons.

Fig S7. T-maze olfactory memory experiment - memory test without electric shock. (A-C) T-maze assays for, the no electrical stimulation *Canton-S* (A), 3 mins interval *Canton-S* (A), 6h interval *Canton-S* (B) and 12h interval *Canton-S* (C). All the interval is after the necessary rest period after training. The numbers on the horizontal line in the figure are MPI, see the METHODS for a detailed description of the fluorescence intensity analysis used in this study.

(D) Neuronal circuitry of the major olfactory neuropils in the *Drosophila* brain. Olfactory sensory neurons (OSN) provide the olfactory input to the first olfactory processing center, the antennal lobe (AL) which consists of ∼ 50 olfactory glomeruli. From the AL, the olfactory information is conveyed via different populations of projection neurons (uniglomerular uPNs and multiglomerular mPNs) to two second-order brain regions, the mushroom body calyx (MBc), and the lateral horn (LH). The MB is composed of intrinsic neurons, called Kenyon cells (KC), which receive direct PN input. The output of the MB to further brain areas is transmitted by a rather small number MB output neurons (MBONs), of which a few also target the LH. The LH is comprised of local neurons (LHLN) and output neurons (LHON) which relay the olfactory information primarily to the SLP (superior lateral protocerebrum), representing the third-order olfactory centers, as well as to the SIP, SMP (superior intermediate/medial protocerebrum), and VLP (ventrolateral protocerebrum). It is conceivable, but has not been proven yet, that the LH sends feedback information to the MB (indicated by the arrow with question mark). In addition, the LH also receives and integrates input from other sensory modalities. LHONs and MBONs have further interactions in third-order neuropils and figure for Scaplen, 2020 ^125^.

(E) Schematic representation of sNPF neurons functioning as accumulatore cells and sNPF-R neurons acting as memory cells in the consolidation process of SELTM

**Movie 1.** Virtual 3D Morphology of YL Neurons. Data from: https://www.virtualflybrain.org/

