## Supplementary figures and images for "Molecular and Neural Circuit Mechanisms Underlying Sexual Experience-dependent Long-Term Memory in *Drosophila*"

### Fig. S1

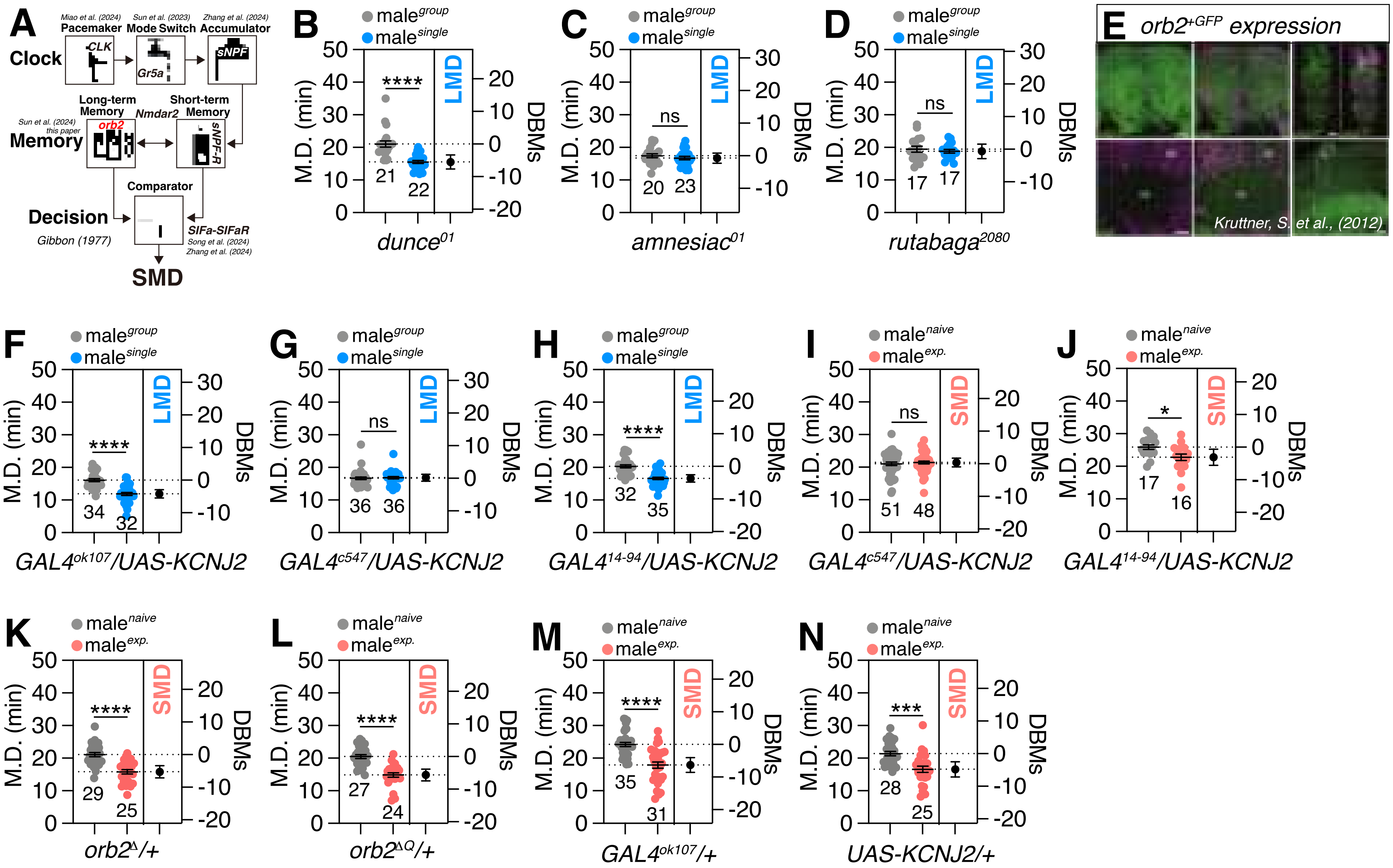


**Memory, Fig.S1**

### Fig. S2

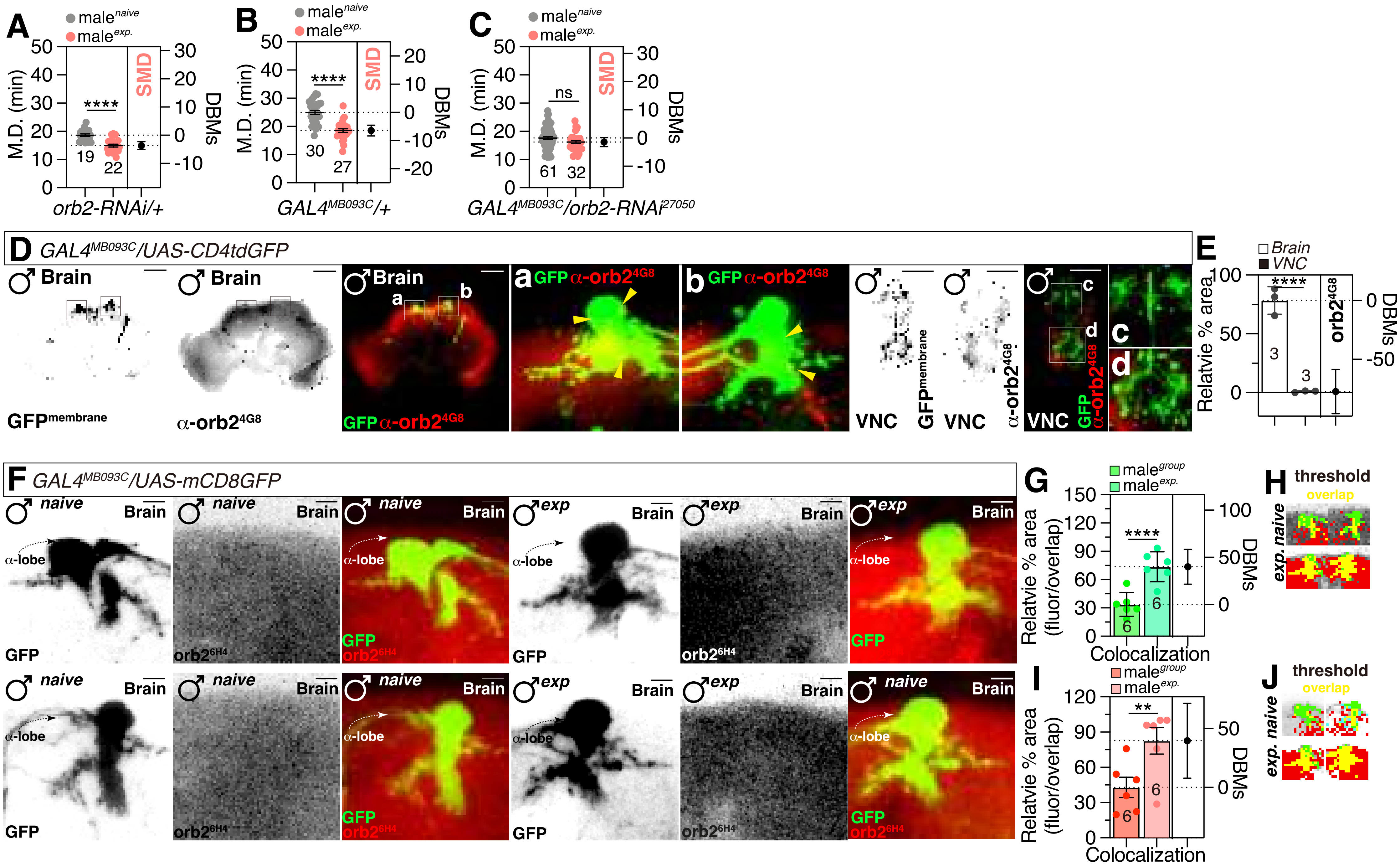


**Memory, Fig.S2**

### Fig. S3

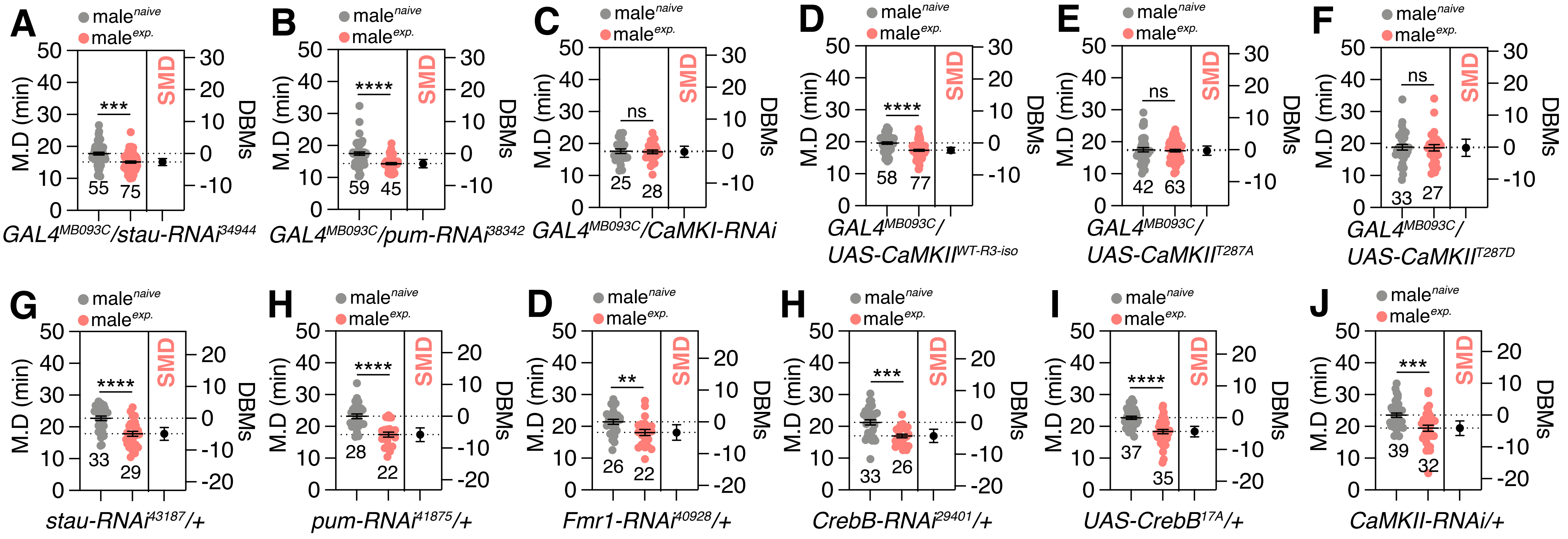


***Memory, Fig.S3***

### Fig. S4

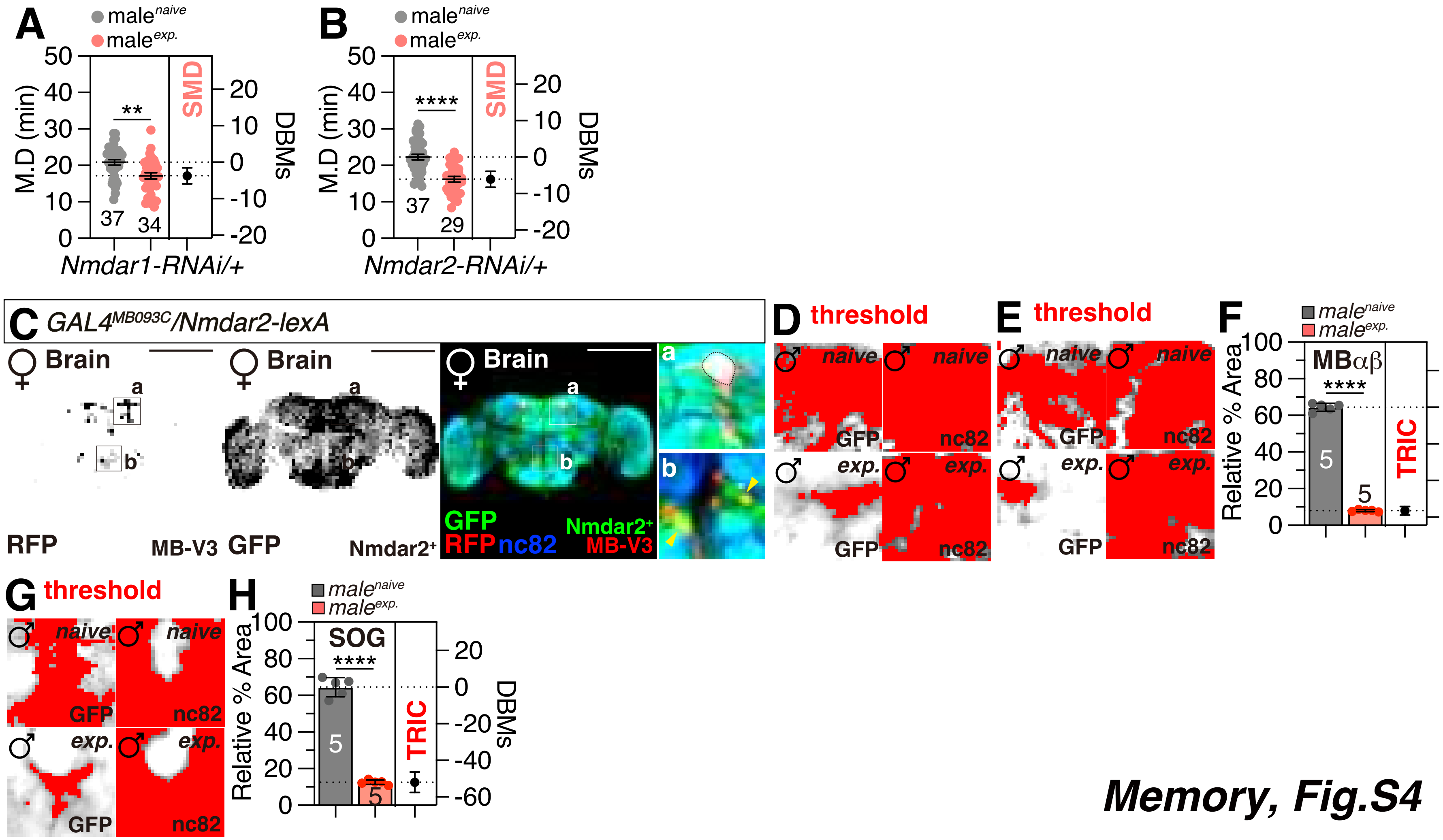


Memory, Fig.S4

### Fig. S5

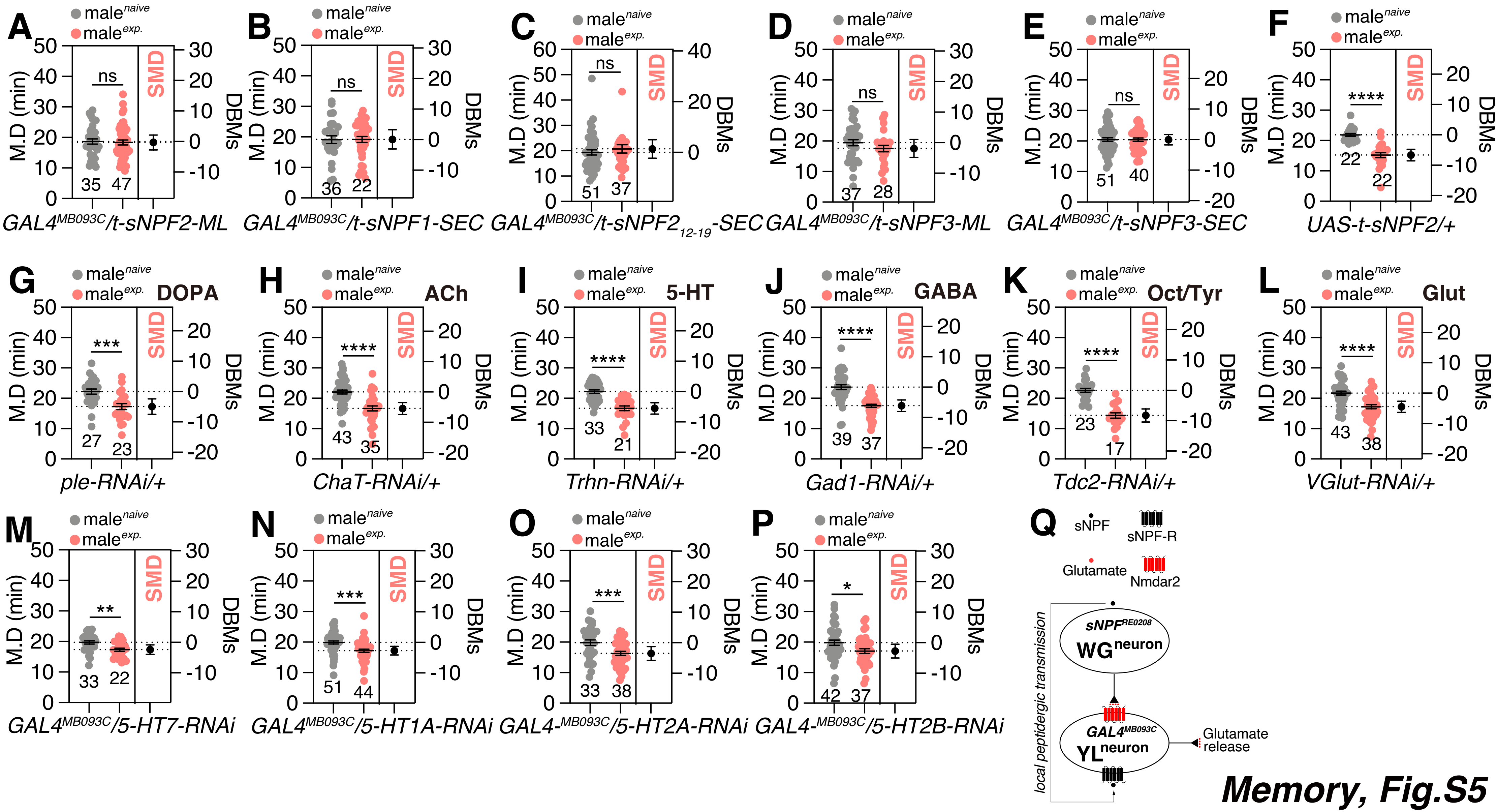


**Memory, Fig.S5**

### Fig. S6

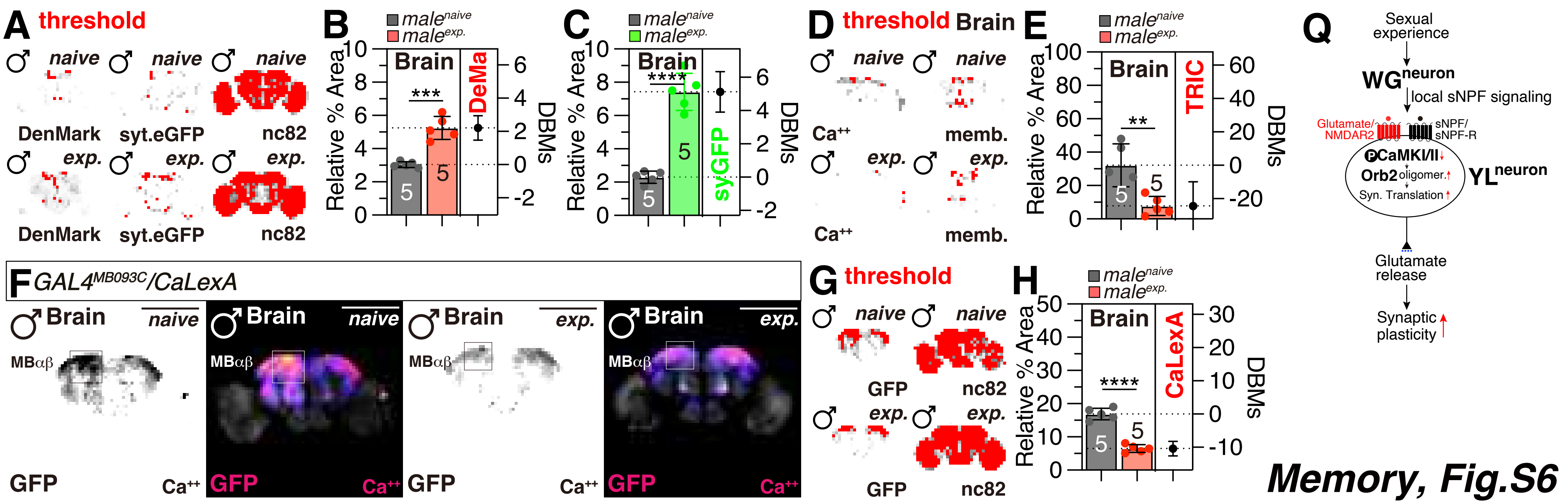


Memory, Fig.S6

### Fig. S7

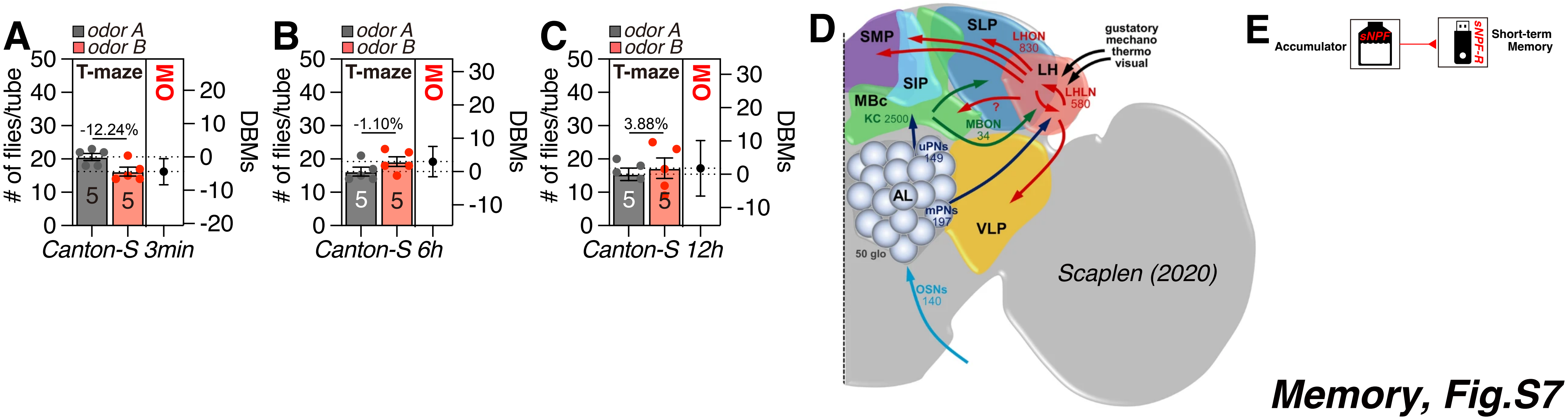


**Memory, Fig.S7**
